# Tackling hysteresis in conformational sampling — how to be forgetful with MEMENTO

**DOI:** 10.1101/2023.01.28.525919

**Authors:** Simon M. Lichtinger, Philip C. Biggin

## Abstract

The structure of proteins has long been recognised to hold the key to understanding and engineering their function, and rapid advances in structural biology (and protein structure prediction) are now supplying researchers with an ever-increasing wealth of structural information. Most of the time, however, structures can only be determined in free energy minima, one at a time. While conformational flexibility may thus be inferred from static end-state structures, their interconversion mechanisms — a central ambition of structural biology — are often beyond the scope of direct experimentation. Given the dynamical nature of the processes in question, many studies have attempted to explore conformational transitions using molecular dynamics (MD). However, ensuring proper convergence and reversibility in the predicted transitions is extremely challenging. In particular, a commonly used technique to map out a path from a starting to a target conformation called targeted MD (tMD) can suffer from starting-state dependence (hysteresis) when combined with techniques such as umbrella sampling (US) to compute the free energy profile of a transition.

Here, we study this problem in detail on conformational changes of increasing complexity. We also present a new, history-independent approach that we term “MEMENTO” (Morphing End states by Modelling Ensembles with iNdependent TOpologies) to generate paths that alleviate hysteresis in the construction of conformational free energy profiles. MEMENTO utilises template-based structure modelling to restore physically reasonable protein conformations based on coordinate interpolation (morphing) as an ensemble of plausible intermediates, from which a smooth path is picked. We compare tMD and MEMENTO on well-characterized test cases (the toy peptide deca-alanine and the enzyme adenylate kinase) before discussing its use in more complicated systems (the kinase P38*α* and the bacterial leucine transporter LeuT). Our work shows that for all but the simplest systems tMD paths should not in general be used to seed umbrella sampling or related techniques, unless the paths are validated by consistent results from biased runs in opposite directions. MEMENTO, on the other hand performs well as a flexible tool to generate intermediate structures for umbrella sampling. We also demonstrate that extended end-state sampling combined with MEMENTO can aid the discovery of collective variables on a case-by-case basis.

**TOC Graphic:** 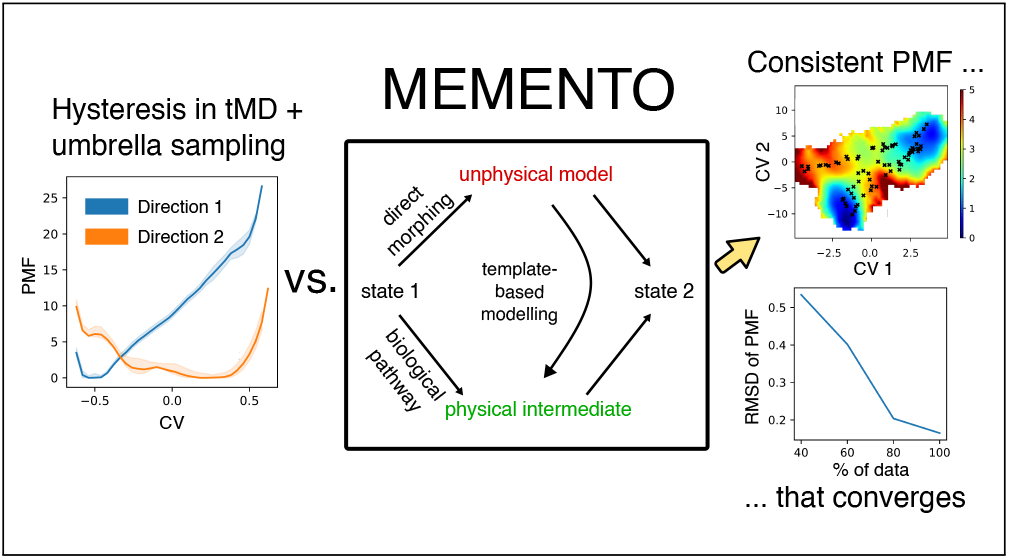

## Introduction

Molecular dynamics (MD) simulations promise to place static protein structures into their dynamical context, which frequently involves large-scale conformational changes.^1^ Thanks to rapid advances in structural biology, one may have information about several conformational states of a given protein available, but the mechanism, energetics and kinetics of their interconversion often remain unknown. This establishes a vital, continuing role for MD in the study of protein structure that comes not without challenges. While tens of microseconds can now be simulated on commercial hardware (rising to hundreds of microseconds with stratification, running multiple boxes in parallel), this is still orders of magnitude shorter than many conformational changes, which often reach into regimes of milliseconds and beyond.^2–4^ Moreover, to accurately describe conformational dynamics at equilibrium, one needs to observe repeated transitions to obtain good statistics. Enhanced sampling methods can help address this issue in various ways: by adaptive spawning of new trajectories, by adjusting potential energy barriers, or by biasing progress along specific reaction coordinates, termed collective variables (CVs).^5^

Several techniques have been developed to implement these strategies. Weighted ensemble methods^6^ use the systematic launching of unbiased MD trajectories to sample along a set of CVs. Temperature-replica exchange MD (REMD)^7^ and accelerated MD (aMD)^8^ are popular strategies to accelerate slow dynamics without reference to collective variables, with modern extensions like replica exchange with solute tempering (REST)^9,10^ and Gaussian accelerated MD (GaMD).^11^ Umbrella sampling (US),^12^ metadynamics^13^ and adaptive biasing force (ABF) sampling,^14^ on the other hand, rely on CVs that capture the relevant slow degrees of freedom (DOFs) of a system. These methods can be highly effective to sample desired conformational changes, however the design of the required CVs is a formidable challenge in itself, often requiring extensive prior knowledge.

With the goal of obtaining a Potential of Mean Force (PMF) — or Free Energy Surface (FES) — of a given conformational change, it is conceptually useful for the majority of the above methods to separate the task into two parts. One first needs to obtain an initial path connecting two known end states, followed by focusing extensive sampling on the vicinity of this path in phase space to gather a PMF. A popular approach is to use targeted MD (tMD)^15^ (also called steered MD) with a CV based on the RMSD to a target state to generate a path, followed by replica-exchange US (REUS) ^16^ and processing with the weighted histogram analysis method (WHAM).^17^ We note here that although time-dependent biasing schemes like metadynamics do not separate path generation and free energy sampling, they still internally construct transition paths by biased sampling and will suffer from similar issues to those described below.

In figure 1 a-b, we present an example of what the tMD + REUS approach may yield in practice. We ran tMD between the DFG-in and DFG-out conformations of P38*α* (details of this system are provided below). In projection onto a simple collective variable (DRMSD = RMSD_(DFG-in)_ − RMSD_(DFG-out)_, where the whole proteins are aligned but the RMSD is calculated along the DFG loop only), both biasing directions appear to produce metastable states in their respective target conformations (figure 1a). Proceeding to REUS, however, reveals substantial starting-state dependence (or hysteresis), illustrated in figure 1b. Generally speaking, when a bias is applied to a CV to move between two conformations, the implicit assumption is that orthogonal degrees of freedom (DOFs) will equilibrate within the available sampling. Where this is not the case, as we show schematically in figure 1c, the result is a PMF skewed to the starting state of path generation.

**Figure 1:**
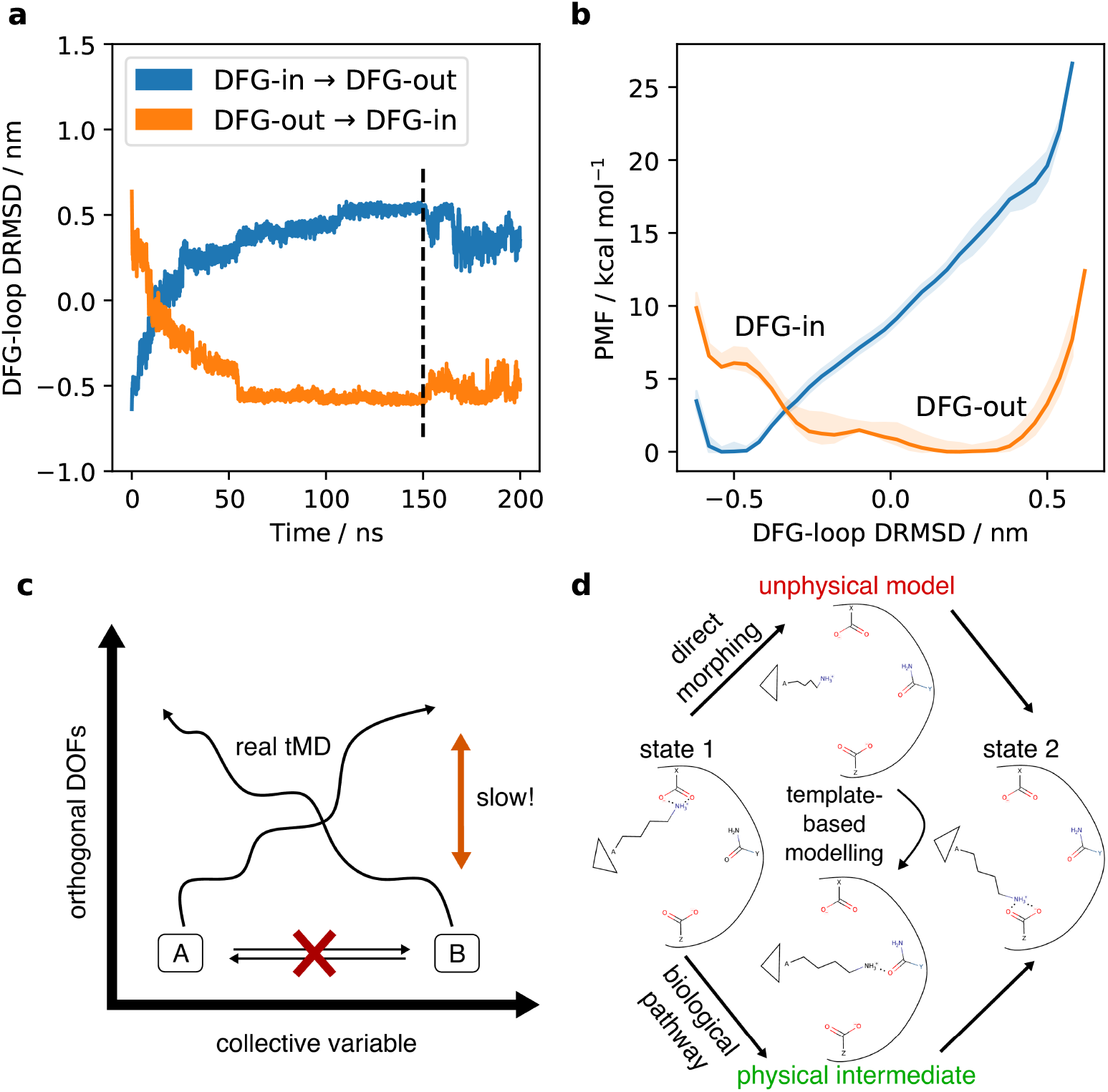
(a) Targeted MD (tMD) between DFG-in and DFG-out conformations of P38*α*. Bi-directional steers on the RMSD to the respective target configuration, projected on the difference of RMSDs (DRMSD). (b) Replica-exchange umbrella sampling (REUS) on the obtained transition paths shows strong hysteresis. (c) Schematic representation of the problem of orthogonal degrees of freedom (DOFs) in tMD. (d) Schematic overview of the MEMENTO procedure, fixing unphysical morphing intermediates by template-based modelling.

There are, broadly speaking, two ways to address this issue. A researcher can attempt to design better CVs that capture all slow movements of the protein by using extensive physical knowledge and restraints (recently demonstrated on a membrane transporter by Meshkin et al.^4^), by combining a larger number of CVs in bias-exchange metadynamics ^18^ or — with some limits on intuitive interpretability — using a host of new machine-learning approaches.^19^ Alternatively, one may wish to construct a path of intermediate structures that is free of hysteresis. In transition path sampling, ^20,21^ an ensemble of transition paths can be constructed from one rare event trajectory, and the string method^22,23^ has been applied with success in transitions as complex as membrane transporter conformational changes. ^24,25^

Since tMD paths like the P38*α* runs we presented above can be highly metastable — to the extent that they appear converged in REUS even with hundreds of nanoseconds of simulation time per window — we decided not to attempt refining paths with MD. Instead, we focus on alternative path-generation algorithms. From various takes on morphing (coordinate interpolation)^26–28^ and rigid-body approximations^29,30^ to elastic network-based models^31,32^ and elaborate analysis–biasing iterative schemes,^33^ the field is ripe with ways to connect protein conformations. Unfortunately, the potential for hysteresis in umbrella sampling often remains understudied because validation of the actual PMFs is rarely undertaken. Furthermore, such validation is hampered by poor code availability and user-friendliness in some of the more complex approaches.

For these reasons, we asked what might be the most conceptually simple, easy-to-implement way to generate paths that can eliminate hysteresis in umbrella sampling. Linear interpolation of coordinates (morphing) is by definition history-free, but the intermediates are unphysical: bonds and angles are distorted and side-chain interactions are disrupted. Many tools exist to fix morphing intermediates (such as RigiMOL in PyMOL^34^ and a panoply of morph servers^27,35–37^) but they have been developed mainly for visual purposes and lack substantial MD validation. On the other hand, the MODELLER package^38^ has been used and improved for decades to prepare starting structures for MD simulations. Here, we find that its algorithm based on satisfying probability distributions for angles and dihedrals together with side-chain interactions is indeed able to fix unphysical morphing intermediates.

This approach for history-free path generation, which we term MEMENTO (Morphing Endstates by Modelling Ensembles with iNdependent TOpologies, illustrated in figure 1d), is validated in this paper on four example systems, the toy peptide deca-alanine, the enzyme ADK, the kinase P38*α* and the membrane-transporter LeuT. At each stage, we rigorously compare our results to what is attainable using tMD alone and find that MEMENTO out-performs tMD on all systems except for deca-alanine, where they behave equally well. We also show how, by combining long end-state sampling with MEMENTO, suitable CVs for umbrella sampling can be developed iteratively. Although we see this research as presenting mostly general findings on how to best sample conformational changes, we also release a documented, tested and user-friendly python implementation of MEMENTO as the path-generation package PyMEMENTO.

## Methods

### Path generation with MEMENTO

#### Basic workflow

To automatically morph protein end states and fix the intermediates with MODELLER, we designed a python package. The source code, documentation and examples are available on github: https://github.com/simonlichtinger/PyMEMENTO. We also provide a static fork of the package taken at the time of publication at https://github.com/bigginlab/PyMEMENTO. Our package uses the MDAnalysis,^39^ numpy,^40^ pandas,^41^ matplotlib^42^ and GromacsWrapper^43^ packages.

In the most basic cases, we perform five key steps.

##### 1. Coordinate interpolation

A linear morph between two protein coordinate files is calculated. If **X**_*i*_ is the coordinate of the i-th atom of a protein, then:

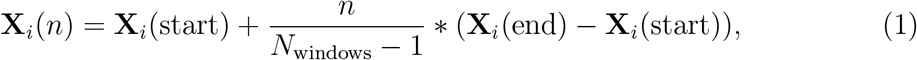

where *n* runs from 0 to *N*_windows_ − 1 and *N*_windows_ is the number of windows one wishes to use for subsequent umbrella sampling. In this work, we find that 24 windows are usually sufficient for good sampling. This is the number of windows used for all examples unless otherwise stated.

##### 2. Template-based modelling

MODELLER^38^ is used to generate *N*_models_ models for each set of coordinates. This fixes the geometry and provides reasonable side-chain orientations and interactions. We achieve this by running model generation with the protein sequence aligned onto itself. In this work, we always use 50 models per intermediate.

##### 3. Finding a smooth path

This ensemble of models at intermediate positions allows potentially for 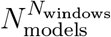 paths between the end states, which is usually an astronomically large number. We therefore attempt to find a smooth path through the model space by running Monte Carlo annealing to minimise 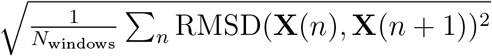, where **X**(*n*) are the heavy-atom coordinates of intermediate *n* out of *N*_windows_. In essence, we minimise the RMSD of RMSDs between neighbouring frames in the path. As a Monte Carlo step, we use the random exchange of a model for another one from the ensemble at one of the intermediates. While it would be difficult to properly converge to a minimum path, here we are merely interested in some appropriately smooth path, so that it is sufficient to take the minimum of 12 independently seeded Monte Carlo runs (which converge to similar results in 10000 steps each).

##### 4. Processing of the results

The frames of the determined path are then subjected to common MD simulation preparation steps: capping of termini (if appropriate), solvating MD boxes, adding ions. We have automated these within the package — for this we rely on the GROMACS simulation engine.^44^ Details are provided below for how ligands and lipids are handled. PyMEMENTO can deal with cubic/orthorhombic and hexagonal boxes as well as the CHARMM and AMBER forcefields.

##### 5. Equilibration

For each of these boxes, in addition to the equilibration protocol referenced under MD simulation details, we run extensive further NPT equilibration with position restraints (1000 kJ mol^−1^) on C*α* atoms, to ensure that all side chains have found suitable interaction partners. For deca-alanine, ADK and P38*α*, we use 10 ns per window, for LeuT and holo-ADK we increase this to 90/100 ns to allow for the membrane and the ligand to equilibrate to a new protein conformation.

From these equilibrated path frames we start our umbrella sampling simulations as detailed in the section on MD simulations. PyMEMENTO provides some additional functionality for setting this up with Gromacs + PLUMED, but the process will fall to the user if a more involved setup (for example 2D umbrella sampling) is required.

#### Adding ligands

It is often instructive to study how a ligand affects the conformational landscape of a protein, and for our ADK and P38*α* systems we have investigated this question with crystallographic ligands. However, for both of these proteins the ligand binding pose is only known for one of the conformational states (and may well not bind tightly to the other). We therefore aligned the end states and manually inserted the holo-state ligand conformation into the apo state, which is not a general approach but in these cases resulted in meta-stable ligand-bound poses after energy minimisation. PyMEMENTO cannot fix internal ligand coordinates, but provided with holo end states thus prepared, it can include the ligand in all intermediate boxes after fractional translation of the centre of mass. We have implemented alternative functionality to linearly morph ligand coordinates and use only energy minimisation to fix them, but we do not employ it in this paper (we foresee it might become important when considering bound ions or crystallographic water in the future).

#### Adding lipids

Our LeuT transporter example case is a membrane protein and therefore needs to be simulated within a membrane box. MODELLER cannot fix lipids and attempting to morph them would likely be futile. Instead, one can provide one of the end states embedded in a lipid membrane. PyMEMENTO uses this pre-equilibrated membrane to embed all other conformational intermediates in lipids in a procedure inspired by the popular inflate-gro script. ^45^ We first stretch the aligned membrane in its plane, followed by cycles of compression and energy minimisation to fit it snugly around the new protein conformation.

### End-state sampling, replicates and two-dimensional CVs

As detailed in the main text, we found great value in running extensive unbiased MD at the end states. Using snapshots from these trajectories to seed MEMENTO, we obtained independent replicates for which we first ran 1D REUS along a naive CV. Where there were significant differences between replicates, we concluded that the end-state sampling had captured a meaningful trend that was propagated through MEMENTO. We therefore concatenated all 1D REUS trajectories (at 5 ns stride for efficiency) and ran principal-components analysis (PCA) on the C*α* positions using the GROMACS implementation. For P38*α*, we did this including only the DFG loop (residues 166-176), and also for the entire protein. The two first PCs obtained in each of these cases were our 2D-CVs for subsequent umbrella sampling.

For LeuT, using the entire protein for PCA, we found that the first PC unsurprisingly correlated very well with the 1D-CV; we used it as the first CV for 2D-REUS (PC 1). However, the contributions from the remaining PCs trailed off rather slowly, requiring 16 PCs to explain >75% of the variance. We thus decided to make a linear combination of PCs 2–16 that would optimally separate out the replicates (under the assumption that this is where the interesting conformational diversity would sit). We built an in-house script that uses scipy^46^ to maximise by differential evolution^47^ an entropy-like metric of distances between MEMENTO path frames:

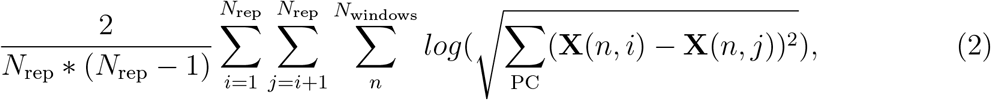

where *N*_rep_ is the number of replicates (here 3) and by **X**(*n, i*) − **X**(*n, j*) we denote the distance between two conformational frames in different replcicates *i* and *j*, evaluated in a projection along a given principal component. The result was termed PC 2 and used as the second CV in 2D-REUS.

While we believe that this approach for iterative CV design can be very useful in combination with MEMENTO, it does not generalise well across systems. Therefore, we did not include it in the PyMEMENTO package which focuses solely on path generation. The design of specific umbrella sampling CVs is left to the user, within the scope of this paper.

### MD simulation details

All simulations in this study were run in GROMACS^44^ in versions 2021.3/2021.4 (the slight version discrepancy is because of different installations on two compute clusters we used). For production simulations, we used the leap-frog integrator with a time-step of 2 fs, the v-rescale thermostat with stochastic term ^48^ with a time constant of 0.5 ps and target temperature of 300 K (310 K for LeuT), the Parrinello-Rahman barostat^49^ with a time constant of 2 ps and target pressure of 1 bar, and a short-range cutoff of 1.2 nm. Where lipids were present, we employed two temperature coupling groups (membrane, protein + water + ions) and a semi-isotropic version of the barostat (x/y, z axes). Example GROMACS *.mdp files are provided with PyMEMENTO.

Before starting production simulations, we energy-minimised and then equilibrated all simulation boxes for 200 ps in the NVT ensemble at a time step of 1 fs, followed by 1 ns in the NPT ensemble at a time step of 2 fs (with the Berendsen barostat), both with C*α* position restraints in place. This was done before the extra MEMENTO equilibrations as described above.

Umbrella sampling simulations were run with PLUMED^50^ in versions 2.7.2/2.7.3 (again, due to different cluster installations). We used bias exchange every 1000 steps (2 ps), and the WHAM algorithm^17^ in the implementation by Alan Grossfield. ^51^ Convergence was assessed — as discussed in the main text and shown in the supplementary figures — by ensuring histogram overlap of neighbouring windows, visualising the PMF using different fractions of the data, and calculating and RMSD between PMFs incorporating successively more data.

Further details are given for the individual systems below. We are making all simulation data available at https://doi.org/10.5281/zenodo.7567883 in the form of key coordinate files, full PLUMED output files with CV projections and WHAM output free energy data. Raw simulation trajectories and various processing scripts will be made available upon reasonable request.

#### Deca-alanine

We used the peptide building functionality of PyMOL^34^ to generate helical and extended conformations of the deca-alanine peptide, which we capped with ACE and NME residues. The simulations were run with the CHARMM 36 forcefield^52^ (version July 2021). The peptide was solvated with 5237 solvent molecules in a cubic box of around 5.5 nm side length. Targeted MD was run starting from the helical state on the C*α* end-to-end distance over 50 ns, with a restraint sliding from 1.446 to 2.666 nm and a force constant of 5000 kJ mol^−1^ nm^−2^. US was done along the same CV, with restraint centers linearly interpolated between the terminal values and a force constant of 1000 kJ mol^−1^ nm^−2^ for 500 ns per window. For the tMD comparison, the starting frames were extracted from the tMD trajectory at even spacing in CV values. The total sampling time expended for deca-alanine was 24 μs.

#### ADK

We obtained structures for ADK in open^53^ and closed^54^ states from the PDB (open: 4AKE, closed: 1AKE — with inhibitor). Since we could simulate all residues (1–214), we did not cap the termini. We solvated the protein with approximately 16,500 solvent molecules (precise number varies between replicates) and a NaCl concentration of 0.15 M in a cubic box of around 8.1 nm side length. The simulations were run using the AMBER ff14.sb forcefield; ^55^ in the holo-state simulations the AP5A inhibitor was parameterised using GAFF2. ^56^ At each of the apo end states, we ran 500 ns of unbiased MD, though the closed state opened spontaneously (as is expected from the PMF), so that our MEMENTO replicates could only be seeded from the initial closed structure and the 0, 250 and 500 ns frames of the unbiased open run. Targeted MD was conducted in 3 replicates in the closed → open direction, and one replicate in the open → closed direction (since asessesing the stability of the resultant conformation was impossible, given that even the native closed state opened spontaneously). We ran these for 150 ns each with a restraint on the C*α*-RMSD to the respective targeted state that linearly increased to 5000 kJ mol^−1^ nm^−2^, followed by 50 ns unbiased equilibration to judge stability of the obtained conformation. For 1D US, we ran three replicates for MEMENTO and tMD starting configurations each (the former prepared as described above, the latter as equally spaced frames from the three opening tMD runs) along the centre-of-mass distance between the ATP-binding LID (residues 123–159) and AMP-binding NMP (residues 31–73) domains. Restraint centers were averaged CV values over the MEMENTO equilibration runs for all boxes, with a force constant of 2000 kJ mol^−1^ nm^−2^. 2D US ran along the LID–CORE (residues 1–30, 74–122 and 160–214) and NMP–CORE distances, with restraint centers again extracted from box equilibrations or the appropriate tMD frames, and a force constant of 1000 kJ mol^−1^ nm^−2^ along each CV. The ‘alternative closed state’ was obtained by clustering (using GROMACS simple linkage with default parameters) MEMENTO-1D-US rep1 window 0 and tMD-1D-US rep1 window 2, as the most occupied cluster in each case. They were incorporated into 2D US by MEMENTO or tMD as described above.

We detail the amount of sampling collected on ADK in table 1.

**Table 1:** Overview of all MD sampling performed on ADK.

| Type of run | Simulation time / ns |
| --- | --- |
| Unbiased MD | 500 * 2 |
| tMD | 200 * 6 |
| 1D US MEMENTO | (259 + 234 + 258) * 24 |
| 1D US tMD | (372 + 240 + 245) * 24 |
| 2D US MEMENTO apo | (147 + 158) * 24 |
| 2D US tMD apo | (158 + 159 + 157 + 250 + 255) * 24 |
| 2D US MEMENTO holo | 164 * 24 |
| **Total** | **76 μs** |

#### P38*α*

We obtained structures for the DFG-in^57^ and DFG-out^58^ states from the PDB (DFG-in: 1P38, DFG-out: 1W83 — with inhibitor). We used PyMOL to mutate two residues in 1P38 to match the sequence of 1W83: H48L and T263A. Note: in the PDB, entry 1P38 was superseded by 5UOJ in 2017, which does not model the DFG loop anymore. However, for consistency with other simulation work we wish to compare our results against,^32^ we still use the 1P38 coordinates, which is validated by the fact that it is a stable conformation in unbiased MD. We simulated residues 4–354 capped with ACE and NME in the AMBER ff14.sb forcefield;^55^ in the holo-state simulations the pyridine-containing L11 inhibitor was parameterised using GAFF2.^56^ We solvated the protein with approximately 27,300 solvent molecules (precise number varies between replicates) and a NaCl concentration of 0.15 M in a cubic box of around 9.6 nm side length. At each of the apo and holo end states (see note in the Adding ligands section), we simulated 500 ns of unbiased MD, the 0, 250 and 500 ns frames of which formed the input structures for the MEMENTO and tMD replicates. For bi-directional tMD, we used the RMSD of all C*α* atoms to the respective target structure as CV, gradually increasing the force constant to 20 000 kJ mol^−1^ nm^−2^ over 400 ns, followed by 100 ns of unbiased relaxation. When proceeding to 1D US, we initially trialed a difference RMSD, DRMSD = RMSD_(DFG-in)_ − RMSD_(DFG-out)_, on all C*α* atoms, but found slightly better behaviour when keeping whole-protein aligning but restricting the RMSD calculation to the DFG loop (resides 166–176). We therefore used this CV for all 1D US, with a force constant of 4000 kJ mol^−1^ nm^−2^ and restraint centers averaged from MEMENTO box equilibrations or extracted from tMD frames (only reps 1 and 3 of the tMD were used for US, since rep 2 did not lead to meta-stable conformations in the vicinity of the target states). We also tried to re-solvate (using the GROMACS solvation tool) the conformations of 1D-US-tMD rep 1 at each intermediate, but stopped our simulations short of the sampling time used in other runs, since we observed no significant difference in PMF. The 2D-US-CVs were derived as discussed in the previous section; we used a force constant of 2 × 10^6^ kJ mol^−1^ on each CV during US (note that the PCs are dimensionless and have small absolute values when output by GROMACS).

We detail the amount of sampling collected on P38*α* in table 2.

**Table 2:** Overview of all MD sampling performed on P38*α*.

| Type of run | Simulation time / ns |
| --- | --- |
| Unbiased MD | 500 * 4 |
| tMD | 500 * 6 |
| 1D US MEMENTO | (150 + 142 + 140) * 24 |
| 1D US tMD (closing) | (148 + 141) * 24 |
| 1D US tMD (opening) | (147 + 130) * 24 |
| 1D US tMD (re-solvation) | (118 + 60) * 24 |
| 2D US MEMENTO apo | (137 + 123 + 101) * 24 |
| 2D US tMD (closing) | (110 + 158) * 24 |
| 2D US tMD (opening) | (179 + 101) * 24 |
| 2D US MEMENTO holo | (132 + 114 + 131) * 24 |
| Total | 64 μs |

#### LeuT

We obtained structures for the inward-facing (IF)^59^ and outwards-facing occluded (OCC)^60^ states from the PDF (IF: 3TT3, OCC: 3F3E), using MODELLER to fix a missing loop in the OCC structure from sequence. We simulate here residues 11–507, with ACE and NME caps, embedded with the CHARMM-GUI membrane builder^61^ in a bilayer of 344 POPE lipids, solvated with approximately 21,500 solvent molecules (precise number varies between replicates) at a NaCL concentration of 0.15 M in an orthorhombic box of around 10.6 * 10.6 * 9.8 nm side lengths. We performed 1 μs of unbiased MD for each end-state, where the 0, 500 and 1000 ns frames served as the starting structures for MEMENTO and tMD replicates. Targeted MD was run on the C*α* atom RMSD to the respective target state with a force constant linearly increasing to 10 000 kJ mol^−1^ nm^−2^ over 250 ns, then held for further 250 ns and followed by unbiased relaxation for 100 ns. 1D US was set up with a distance RMSD CV, DRMSD = RMSD_(IF)_ − RMSD_(OCC)_, calculated on all C*α* atoms. The force constant was 20 000 kJ mol^−1^ nm^−2^ and restraint centers were averaged from MEMENTO box equilibrations. The derivation of 2D CVs is described in the section above; we used a force constant of 5×10^6^ kJ mol^−1^ along each CV. We used the same MEMENTO starting frames as for 1D US, and tMD frames extracted at equal spacing (only reps 1 and 2 of the OCC → IF tMD were used for US, since the others did not lead to meta-stable conformations in the vicinity of the target states). To improve histogram overlap, we included additional windows in our 2D-US: (1) a MEMENTO run between the starting and final conformations of the IF unbiased run for the 2D-MEMENTO-FES, (2) frames extracted from the IF unbiased run at equal CV spacing for the 2D-tMD-FES, (3) a MEMENTO run with 16 windows between windows 10 and 12 coordinates of MEMENTO rep 3, with higher force constant 2 × 10^7^ kJ mol^−1^, for the 2D-MEMENTO-FES (4) a MEMENTO run with 16 windows between windows 9 and 12 coordinates of MEMENTO rep 2, with higher force constant 2 × 10^7^ kJ mol^−1^, for the 2D-MEMENTO-FES.

We detail the amount of sampling collected on LEUT in table 3.

**Table 3:** Overview of all MD sampling performed on LEUT.

| Type of run | Simulation time / ns |
| --- | --- |
| Unbiased MD | 1000 * 2 |
| tMD | 600 * 6 |
| 1D US MEMENTO | (156 + 155 + 157) * 24 |
| 2D US MEMENTO | (495 + 514 + 517) * 24 |
| 2D US MEMENTO extra | 510 * 24 + (523 + 499) * 16 |
| 2D US tMD | (507 + 511) * 24 |
| 2D US tMD extra | 500 * 24 |
| **Total** | 118 μs |

## Results

### Deca-alanine

We first sought to verify that MEMENTO can indeed fix unphysical morphs without introducing artifacts. Therefore, we applied the method to a simple toy model: deca-alanine in water (see Methods for details on simulation setup). This peptide undergoes a transition between *α*-helical folded and unfolded conformations that has a significant energetic and entropic barrier, but has also been successfully described with US in the literature.^62^ Figure 2 a-b and the step-through supplementary video 1 illustrate how reasonable bond lengths, angles and dihedrals are reconstructed by MEMENTO. We ran one-dimensional REUS from these conformations, using the end-to-end distance of the peptide as a CV, and found a sharp free energy minimum at the helical state as well as a broad minimum at an ensemble of extended states (figure 2c). We note that the shape of this PMF is compatible the literature.

**Figure 2:**
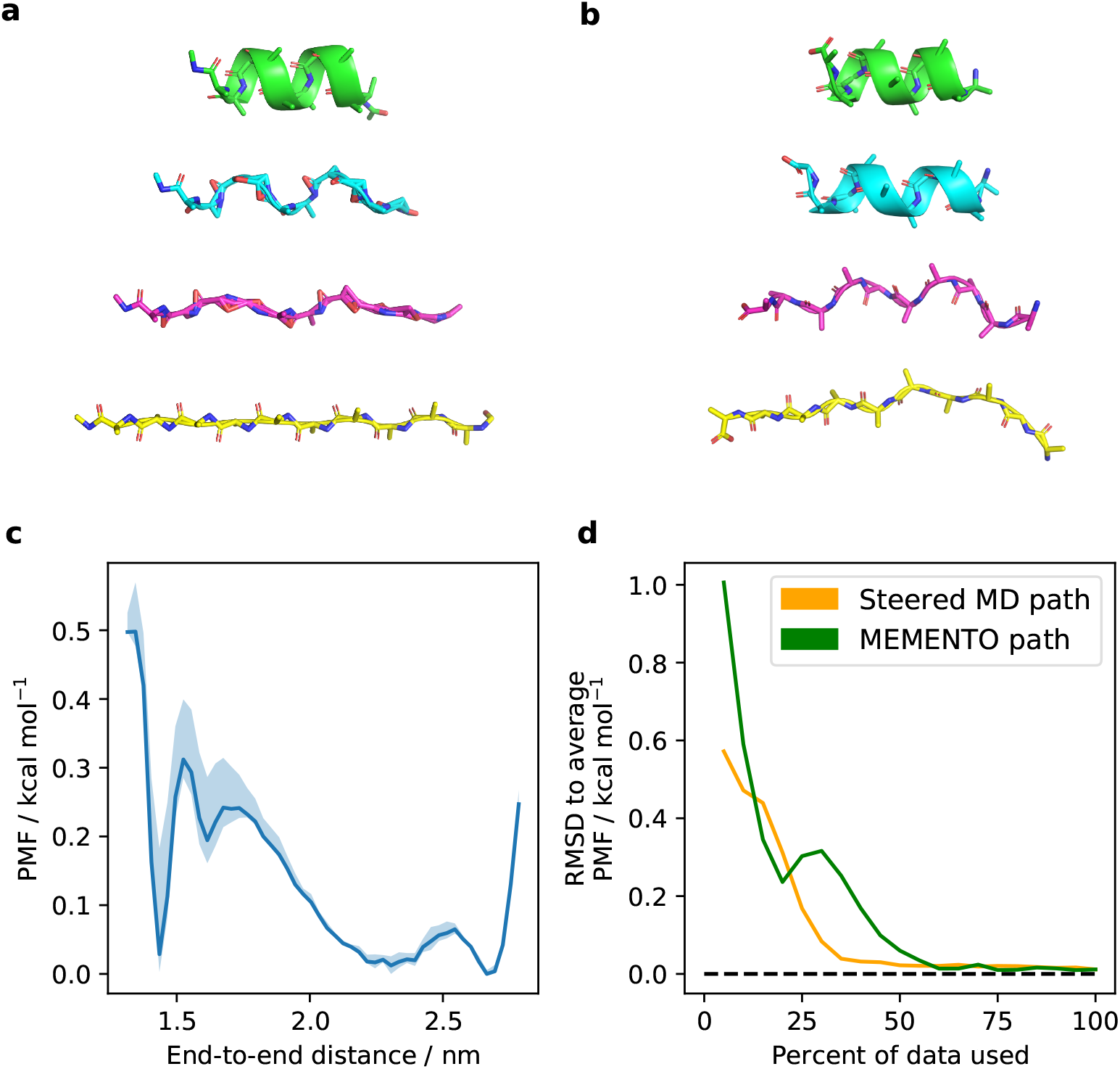
(a) MEMENTO windows 0, 7, 15 and 23 after linear morphing, showing unphysical geometry in the intermediates. (b) The same windows after processing and path-picking. Unphysical geometry is now fixed. (c) PMF of deca-alanine unfolding in water, sampled by 1D REUS along the end-to-end distance. The shaded area is the range of PMFs observed when taking only the first 60%, the last 60% and the full sampling, which gives an indication of error and convergence. (d) MEMENTO and steered MD paths converge roughly equally fast, as measured by the RMSD of PMFs to the final PMF (averaged between MEMENTO and steered MD) when including increasing chunks of sampling.

To further compare MEMENTO with existing methods, we also ran targeted MD along the end-to-end distance CV to open up the helical state (figure S1a). REUS from snapshots along this trajectory yielded a PMF virtually identical to MEMENTO (figure S1b), which also converges well on a similar time scale (figures 2d, S1 c-f). We conclude that MEMENTO performance is on a par with tMD for simple systems, and that it produces high-quality molecular conformers that behave well in US.

### Adenylate kinase (ADK)

While the previous example has established that MEMENTO does not degrade sampling of a system where sampling issues have no impact within the available timescale, we begin to see its benefits when moving to more complex systems. Adenylate kinase (ADK), an enzyme that catalyses the interconversion of adenosine phosphates, is known to exhibit a large-scale conformational transition between open^53^ and closed^54^ states (figure 3a). This conformational change may be described either by a varying distance between the LID and NMP domains, or by the distance of LID and NMP domains from the protein CORE. For many years ADK has served as a benchmark system for computational research on conformational changes and enhanced sampling, some of which we summarise in Table 4. While different studies have obtained substantially different results with various methodologies, the clear literature consensus of atomistic studies is that apo ADK favours the open state while the presence of substrate or inhibitor stabilises the closed conformation. This is also consistent with the crystallographic data and mechanistic intuition.

**Table 4:**
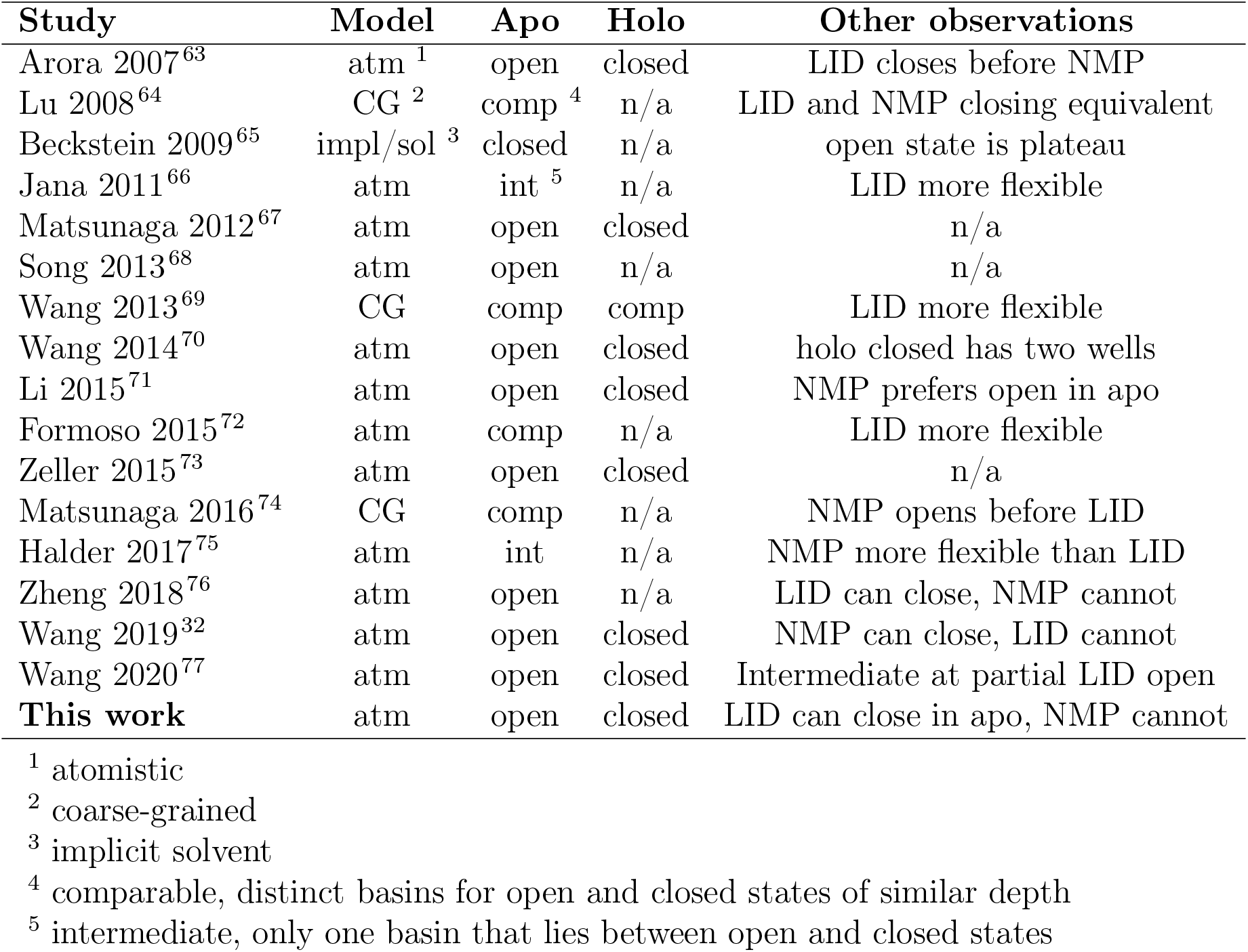
Comparison of our results to published PMFs of the ADK conformational change. We indicate the type of computational model used and which state was found to be favoured in the apo and holo proteins. Our results agree with the consensus of the atomistic studies, while coarse-grained models appear to not capture the conformational changes well.

**Figure 3:**
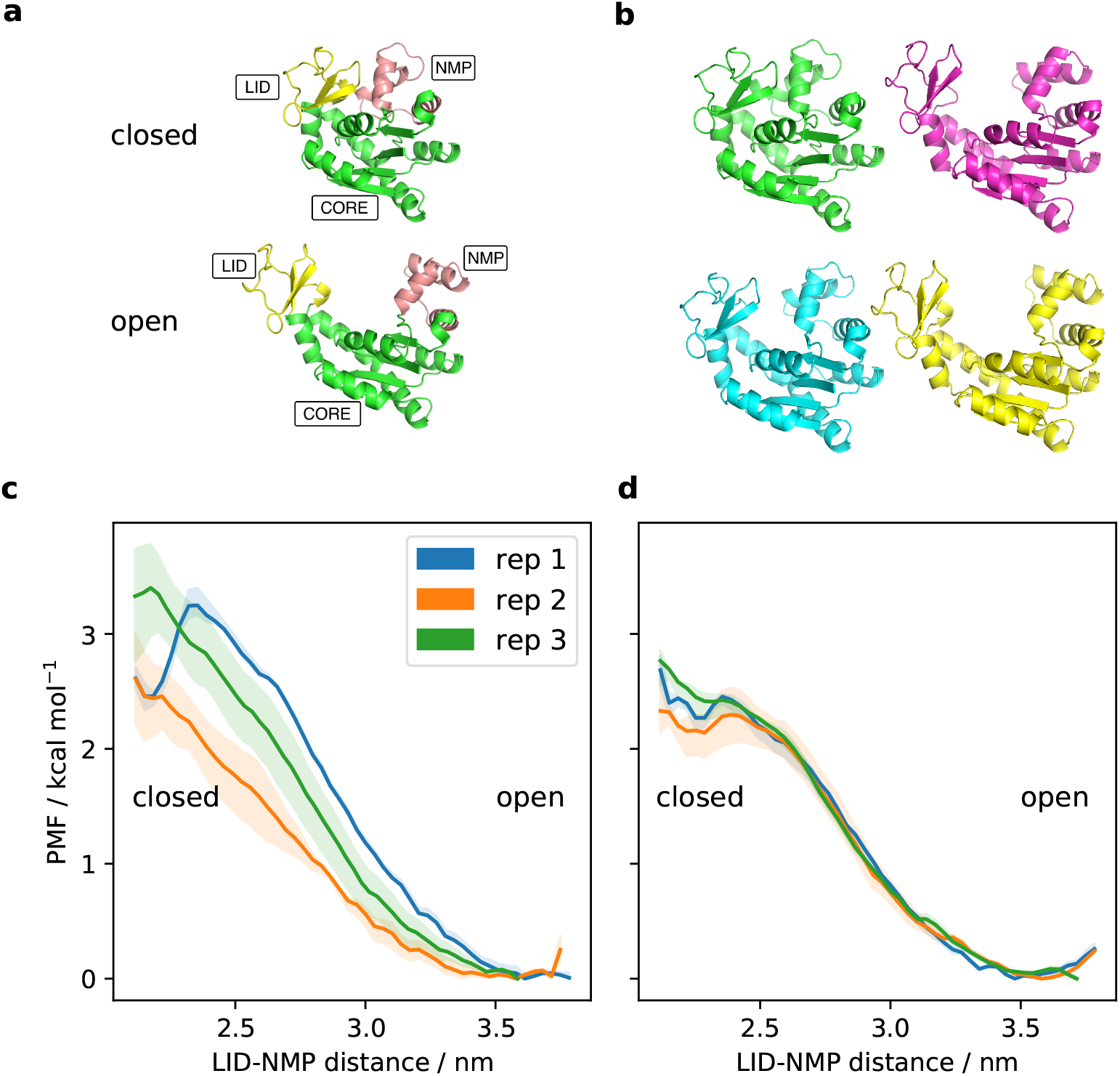
(a) An overview of the domain motions in the ADK open closed conformational change. (b) Representations of MEMENTO ADK intermediates 0, 7, 15 and 23, displaying the required domain motion. (c)–(d) PMFs from 1D-REUS of the ADK conformational change along the LID–NMP CV, with paths from (c) MEMENTO and (d) tMD. Shaded areas are the ranges of PMFs observed when taking only the first 60%, the last 60% and the full sampling.

We applied the MEMENTO procedure to ADK (intermediates are shown in figure 3b and step-through supplementary videos 2–3), where we performed independent replicates by equilibration of the open state in unbiased MD (see Methods for details, closed-state equilibration could not be used to seed replicates due to spontaneous opening in MD). We also ran three replicates of tMD based on a C*α*-RMSD CV in the closed → open direction, and one replicate in the open → closed direction (figure S2 a-b; since the closed state opened in unbiased MD, we had no measure for how successful the open → closed tMD direction might be — thus we did not pursue it further). Using the LID–NMP domain distance as a simple one-dimensional CV, we computed PMFs for our three MEMENTO and tMD replicates (figure 3 c–d, convergence analysis in figure S2 c–f).

As expected, the open state was favoured over the closed state by a roughly similar free energy differences in all replicates. However, some differences between replicates remained, which prompted us to investigate what a relevant orthogonal DOF may be in this case. By clustering windows 0 (MEMENTO) and 2 (tMD) of the respective first replicates, we found a meta-stable alternative closed state (figure 4a), in which the LID is closed on the CORE, but the NMP domain remains open. On inspection, this conformation can be rationalised, because a number of positively charged residues that coordinate the highly negatively charged substrate in holo ADK (see figure 4c) can be engaged in alternative salt-bridges in the absence of ligand. By connecting this alternative closed state to the open conformation with another MEMENTO path, we captured a 2D PMF along the LID–CORE and NMP–CORE distances as CVs (a combination used previously in the literature^32^), shown in figure 4b, that converges very well (figure S3). In a separate set of simulations including the crystallographic inhibitor AP5A, we likewise achieved a converged PMF — this time favouring the closed protein conformation (figure 4d). Consistent with the majority literature conclusions, taken together these PMFs show how the crystallographic closed state is unstable in apo ADK, but also that the LID domain has significant flexibility in the open state — driven by non-native salt-bridge interactions that compensate partially for the lack of substrate. By contrast, holo ADK prefers the closed state, where a number of positively charged residues form salt bridges with the ligand.

**Figure 4:**
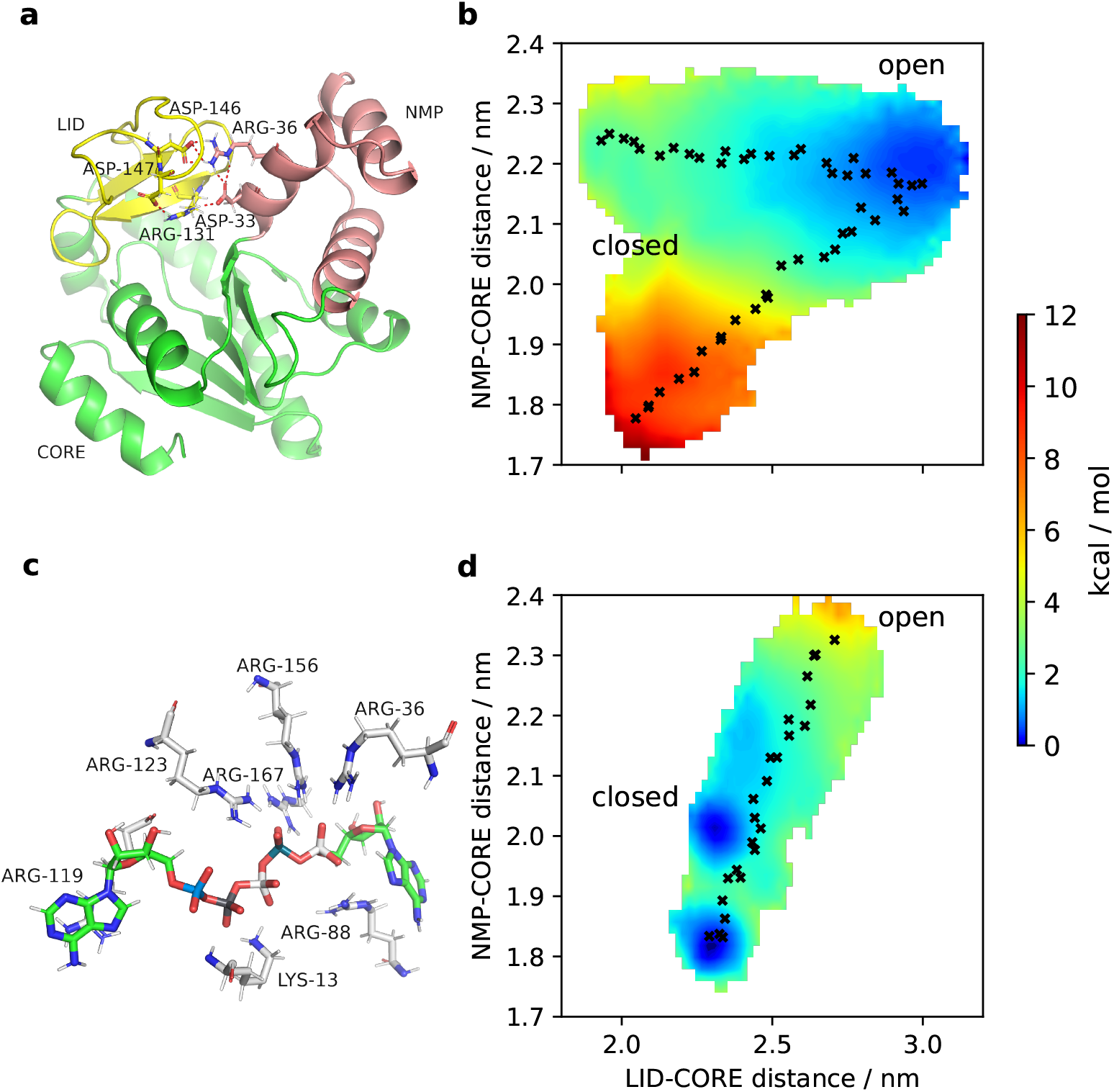
(a) An alternative closed state identified by clustering from 1D-REUS with MEMENTO paths. The LID domain is closed while the NMP domain remains open. (b) PMF from 2D-REUS along the LID–CORE and NMP–CORE CVs, with MEMENTO paths connecting the closed, open and alternative closed states. Crosses indicate the REUS window starting frames. (c) AP5A inhibitor binding pose in the 1AKE crystal structure, illustrating how the negative ligand charge is accommodated by multiple positively charged protein residues. (d) PMF from 2D-REUS in the presence of AP5A, displaying a switch of the conformational preference to the closed state.

We also investigated whether a similar PMF can be recapitulated using only tMD paths. When we projected our tMD runs onto the 2D-CV space, we discovered that the closed → open runs opened the LID before the NMP domain, and the open → closed run closes the LID domain before the NMP domain (figure S4a). In light of our MEMENTO results regarding LID flexibility this is encouraging, however an attempt to compute a 2D-PMF by REUS (figure S4b) converged poorly (figure S4 c–d), even if the rough shape of the PMF was consistent with our previous results. Furthermore, we did not succeed in incorporating the alternative closed state explicitly into the PMF, because REUS from bi-directional tMD between the open and alternative closed states showed significant hysteresis (figure S4 e–f). We therefore conclude that MEMENTO not only produces results in line with expectations for ADK, but also that there is a tangible advantage over tMD for path generation in this case. The example further highlights how MEMENTO can help identify orthogonal DOFs and thus design better CVs through the possibility of incorporating extensive end-state sampling to propagate conformational diversity that would not otherwise be feasible to achieve across many stratified windows. There is also full flexibility to add extra paths to an existing PMF, which is useful if one does discover relevant alternative conformations.

### P38*α*

To provide a second example of a conformational change in a globular protein (that is not as much of a canonical example in the enhanced sampling field as ADK, but for which previous work still exists) we next focused our efforts on P38*α*, a mitogen-activated protein kinase with roles in heart physiology and implications in heart disease. ^78^ P38*α* undergoes a conformational change in its DFG loop, where the DFG-in conformation^57^ is active, while the DFG-out state^58^ is inactive (illustrated in figure 5a). It is known from NMR^79^ and EPR^80^ studies that the apo P38*α* displays a conformational equilibrium between DFG-in and DFG-out geometries, and that type 2 inhibitors (which bind to the DFG-out conformation) reduce the accessibility of the active DFG-in conformation. Several computational studies have broadly come to the same conclusions using various enhanced sampling methods,^32,70,81^ although inhibitors — where they were simulated — are not found to abolish completely the (meta-)stability of a DFG-in-like state, but merely raise it in energy relative to DFG-out.

**Figure 5:**
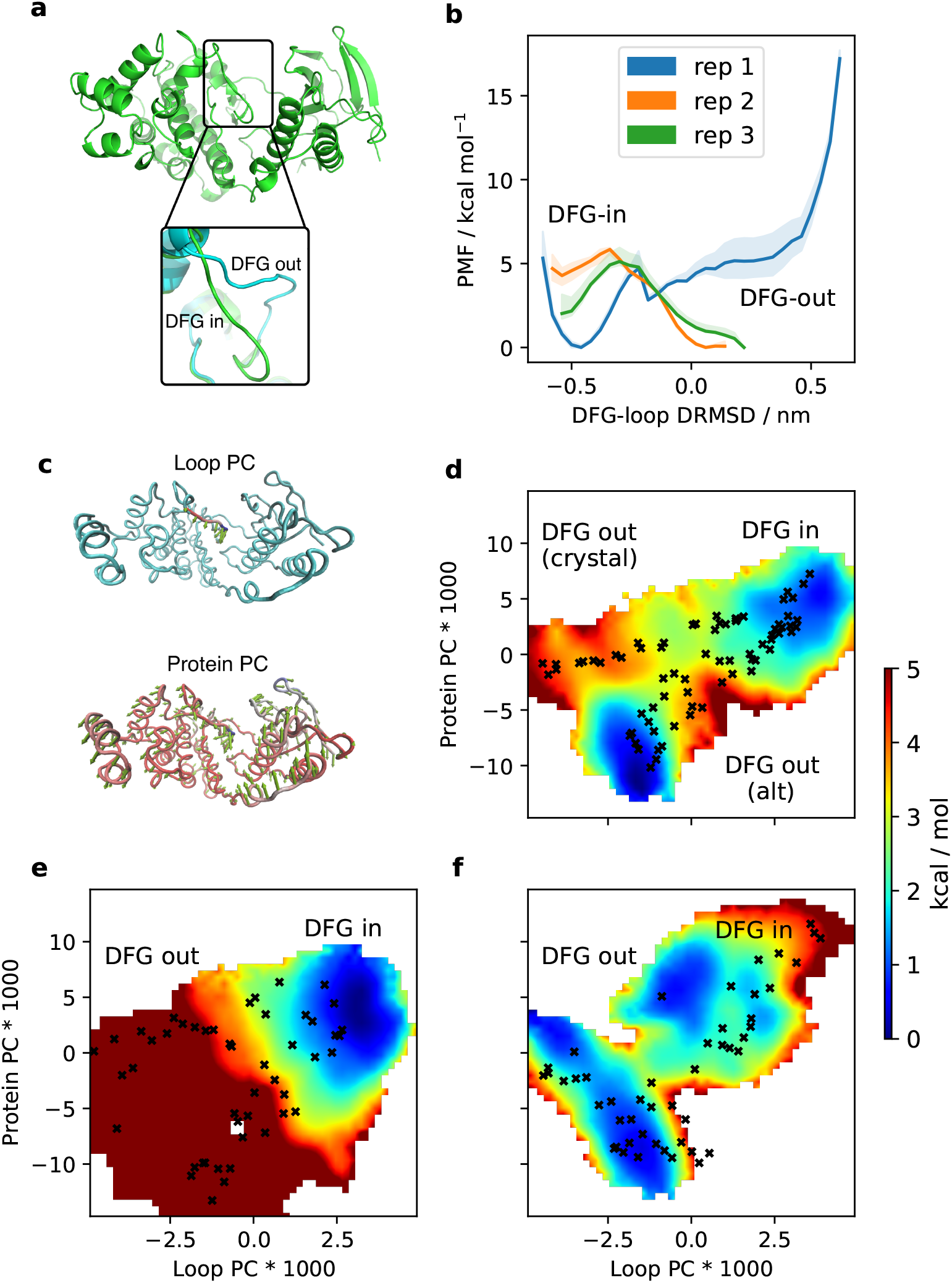
(a) Overview of the P38*α* system, highlighting DFG-in and DFG-out states. (b) 1D-REUS along the DRMSD CV from MEMENTO paths, showing big differences between replicates. Shaded area is the range of PMFs observed when taking only the first 60%, the last 60% and the full sampling. (c) PCA results, showing the two CVs obtained for 2D-REUS sampling. (d) 2D-REUS along PCA-CVs with MEMENTO paths. Crosses indicate the REUS window starting frames. (e)–(f) 2D-REUS from tMD paths, in the (e) DFG-in → DFG-out and (f) DFG-out → DFG-in directions.

As for ADK, we first simulated 500 ns of unbiased equilibrations at all combinations of DFG-in/out and apo/holo. In the introductory figure 1 a-b, we have already demonstrated how tMD suffers from large hysteresis in 1D-REUS along a difference RMSD calculated for the DFG-loop only (DRMSD). To see how a history-independent path would perform in comparison, we ran three MEMENTO replicates (step-through supplementary videos 4–5), followed by 1D-REUS along the DRMSD (figure 5b). While the results already look more reasonable than tMD, the substantial differences between replicates we found led us to hypothesise that our end-state sampling included significant conformational relaxation which was propagated through MEMENTO, but was of too large a timescale to be equilibrated in each REUS window. To find a set of CVs which includes this slow DOF, we opted to extract relevant information from the 1D-REUS sampling we had already accumulated, in an iterative fashion via principal component analysis (PCA; see Methods for more details). By including the DFG loop only or the whole protein in the PCA, we hoped to obtain principal components (PCs) that separate the DFG-loop motion from a superimposed protein conformational bias (figure 5c and supplementary videos 6–7 illustrate the highest contributing PCs in each case, which we used as CVs in subsequent simulations). The loop PC expectedly covers a motion that would interconvert DFG-in and DFG-out loop conformations. The Protein PC also carries an element of DFG-loop motion, but in conjunction with a twist in the protein along the thin hinge that lies under the DFG loop.

Subsequent 2D-REUS we ran along the same MEMENTO paths with the new CVs produced a PMF that was highly instructive about the processes at work (figure 5d). Projecting our initial apo-P38*α* unbiased MD (that started from the crystallographic DFG-out conformation) onto this PMF (figure S5), one can clearly see how the protein assumes an alternative, less extreme DFG-out geometry on removal of the crystallographic ligand. Replicates 2 and 3 therefore start from this free energy basin, which explains why these replicates showed an increasing bias towards DFG-out in 1D-REUS, and also why replicate 1 converged poorly in the DFG-out region. Because the relaxation movement is now included in the PMF, convergence stands much improved and is no longer worse in some regions of the PMF than others (figure S6).

With a working set of CVs at hand, we further wished to investigate whether the 1D-REUS hysteresis with tMD paths we observed was only caused by our poor initial 1D-CV guess, or whether the path itself exhibited starting-state dependence. We therefore projected our tMD replicates onto the 2D-PMF (figure S7a) and found that replicates 1 and 3 appeared to lead to meta-stable states in the vicinity of their respective targets. However, in 2D-REUS, the paths still showed significant bias towards the tMD starting states (figure 5 e–f). Within similar amounts of sampling as used for MEMENTO, this is likely not a convergence issue (figure S7 c–f), although the DFG-out → DFG-in tMD direction has some issues with REUS histogram overlap leading to slower convergence. We also investigated whether the hysteresis might be predominantly due to slow solvent DOFs that could be addressed by re-solvating the tMD intermediates, but found that this does little to improve the situation (figure S7b). In conclusion, while better CVs may well theoretically capture those DOFs that produce hysteresis in tMD for P38*α*, they are not as simple to obtain as with MEMENTO (and we have not managed to develop any here).

Finally, we wished to explore the role of the crystallographic type 2 inhibitor that was bound in the DFG-out structure^58^ (figure S8a). Using the same 2D-CVs as before, we obtained a converged PMF (figure S8 b–d) that showed a free-energy basin near the DFG-out crystal structure, in contrast to the apo state. However, the PMF displays low free energy barriers to conformations that register as DFG-in to our CVs, which contradicts the experimental evidence. Noting that other MD studies (using 1D US)^32,70^ also found DFG-in conformations to be accessible in holo P38*α*, we judge that this may either be an issue with MD models of this system, or that what we observe as DFG-in-like in our simulations may not be experimentally relevant DFG-in conformations, thus hinting at another CV problem. To our knowledge, this issue has not been addressed before — and it would be worthwhile to do so in the future — however we see this as beyond the scope of the present study.

### Leucine transporter (LeuT)

Thus far, our validation examples were all soluble proteins. However, conformational dynamics of other types of systems such as membrane proteins also play a crucial role in biology. For example, the solute carrier (SLC) superfamily encompasses 65 families of more than 450 transport proteins, which facilitate the movement of substrates ranging in size from protons to steroids and heme across the cell membrane.^82^ These transporters function by exposing a substrate binding site in turn to the intra- and extracellular medium, in what is termed an alternating access mechanism. Among the SLC transporters, many structurally diverse variants have been described, which operate by the rocker-switch, rocking bundle and elevator mechanisms. ^83,84^ A dynamical reprsentation of these conformational changes is of great pharmacological interest since many drugs are carried by SLC transporters, and may help design better drug delivery approaches in the future.

LeuT — a bacterial sodium-coupled leucine transporter — is an archaetypal SLC transporter with a rocking bundle mechanism (the folds observed in the SLC 6, 7, 11, 12 and 38 families are its namesakes). Therefore, understanding the LeuT conformational changes is likely to facilitate insight into the transport of various amino acids, neurotransmitters and inorganic ions. Here, we apply MEMENTO and tMD to the conformational change between the inwards-facing (IF)^59^ and outwards-facing occluded (OCC)^60^ conformations (figure 6a).

**Figure 6:**
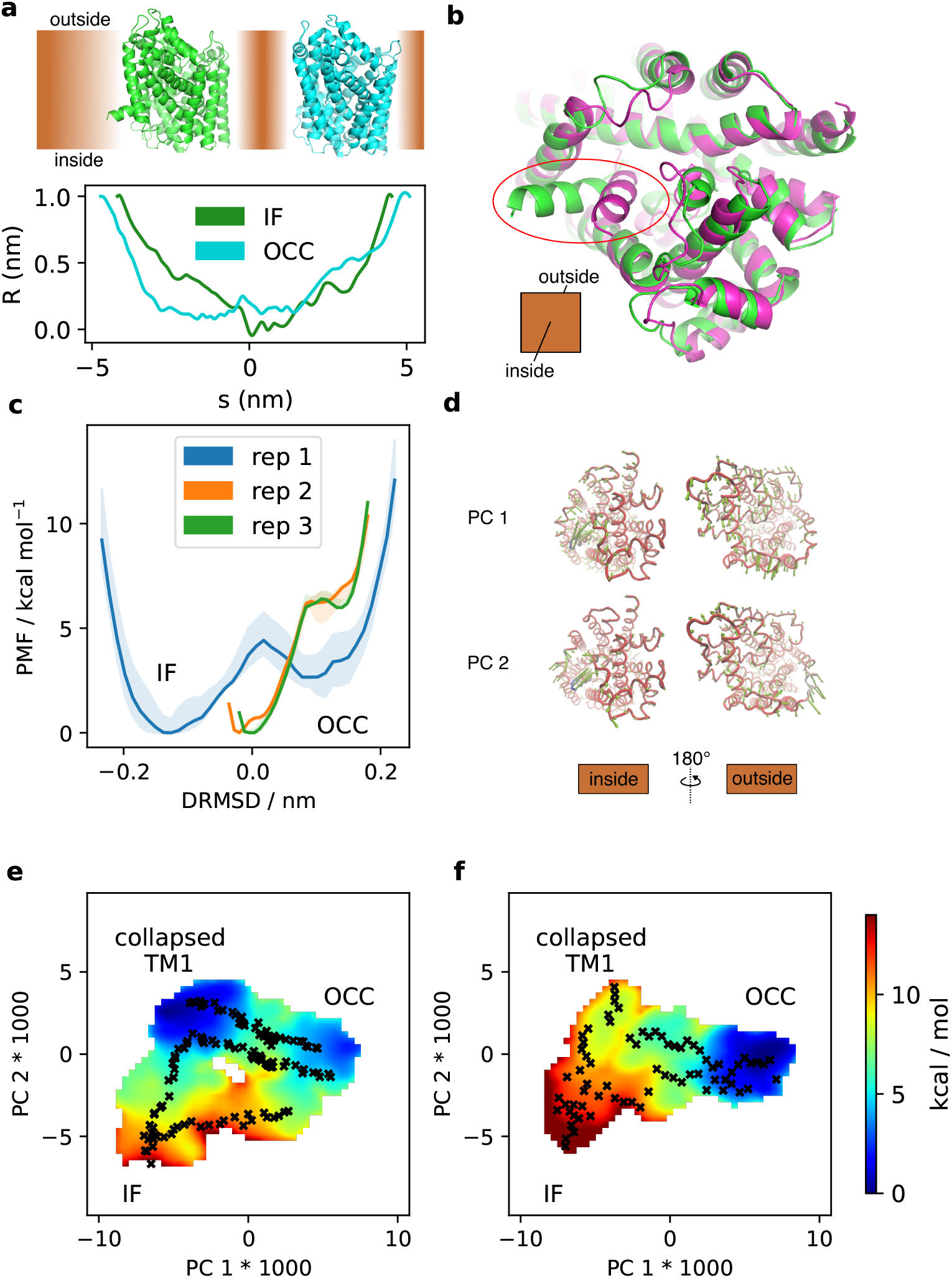
(a) Overview of the IF and OCC structures of LeuT, and pore radius profiles calculated with CHAP.^88^ (b) Illustration of the closing motion of TM1, before (green) and after (pink) 1 μs unbiased MD. (c) 1D-REUS along the DRMSD CV from MEMENTO paths, showing big differences between replicates. Shaded area is the range of PMFs observed when taking only the first 60%, the last 60% and the full sampling. (d) PCA results, showing the two CVs obtained for further sampling. (e)–(f) 2D-REUS from (e) MEMENTO and (f) tMD paths. Crosses indicate the REUS window starting frames.

As in the previous examples, we first ran long (here 1 μs) unbiased MD on lipid-embedded boxes of these two conformations (both as apo protein). While the OCC state is stable in MD, we noticed that in the IF simulations, transmembrane helix 1 (TM1) collapsed at its intracellular end to an orientation that takes it closer to the opposing bundle — that is, it partially closed spontaneously (figure 6b). Propagated through MEMENTO paths (step-through supplementary videos 8–11) and 1D-REUS along a DRMSD CV (based on the original IF and OCC coordinates), this lead to big differences between replicates. Notably, in replicates 2 and 3 (the starting states of which were ‘collapsed TM1’), the PMF is not sampled near the original IF structure (figure 6c). Since we wished to determine the energetics of the full connecting path between the IF and OCC structures, we once again set out to construct 2D-CVs using PCA based on the 1D-REUS trajectories. The details of this procedure are described in the Methods; in short, we found that too many PCs were required to explain a substantial proportion of the variance to be directly useful in US. Therefore, we took PC 1, the highest contributing direction, as one CV, while constructing the second CV as a linear combination between the next 15 PCs that separates the replicate MEMENTO paths best (we termed this CV ‘PC 2’ for simplicity). These CVs are global and complex, but roughly speaking PC 1 covers the conformational changes occuring both at the intra- and extracellular sides between IF and OCC, while PC 2 focuses more on the TM 1 movement with less contributions at the extracellular face (figure 6d and supplementary videos 12–15).

Equipped with these CVs, we proceeded to estimating the 2D-PMF of the overall conformational change. In our first attempts, we experienced issues with undersampling the IF ↔ ‘collapsed TM1’ region (unsurprisingly, since it was not originally included as an explicit path) as well as parts of the OCC ↔ collapsed TM1 transition region (it appears that 24 windows were not quite enough for this conformational change). We therefore used MEMENTO again to supplement our PMF with extra paths between the crystallographic and ‘collapsed TM1’ IF states, as well as additional 16 windows along the replicate 2–3 paths within the transition region. The final PMF is shown in figure 6e and converges reasonably well (figure S9), though somewhat worse than previous examples even with 500 ns / window of sampling. The PMF reproduces the observation from unbiased MD that the IF state is not stable in the simulated conditions but collapses into a broad basin of an inward-facing, partially occluded state. There is then a second free energy barrier between this basin and the crystallographic outwards-facing OCC state, which corresponds to the rocking bundle motion. Since this barrier is significantly smaller however than for the direct IF ↔ OCC interconversion, our simulations suggest a preferred sequential mechanism for the overall conformational change proceeding via first closing TM1 before engaging in the rocking bundle movement.

This PMF agrees well with previous data published on LeuT. Gur et al. ^85^ simulated a set of unbiased MD trajectories, starting from crystal structures and paths obtained using an anisotropic network model-based biasing scheme termed coMD. They found two free energy basins of inward-facing and outward-facing conformations that interconvert most favourably via occluded states (although obtaining sufficient sampling in the transition region was challenging). They also found that the intracellular side of TM1 tends to partially close up during their situations. This observation had also been made in previous experimental^86^ and computational^87^ investigations, and is reproduced in this work in the ‘collapsed TM1’ state. The literature therefore supports the shape of the PMF we have obtained for apo LeuT. Moreover, to our knowledge this study contributes the most extensive sampling of LeuT IF ↔ OCC conformational dynamics expended to date, therefore corroborating earlier results. However, investigating the role of substrate and coupled sodium ions in the transport cycle — as other studies have attempted — was beyond the scope of the present work, since we are chiefly concerned with sampling methodology here.

In order to compare our MEMENTO 2D-PMF to what would be achievable with tMD for path generation, we ran three replicates of long (500 ns) tMD, with initial and target states set to the same frames of unbiased MD as we used for MEMENTO above (figure S10). We found that only tMD OCC → IF replicates 1–2 and no IF → OCC runs reached meta-stable conformations near their target states. When we calculated a 2D-PMF along these paths, supplemented with extra frames from the apo IF unbiased run (in place of the extra MEMENTO path used above to close the gap between IF and ‘collapsed TM1’), we found a strong bias towards the OCC state (figure 6f) and convergence worse than with MEMENTO paths (figure S11). The fact that the windows from unbiased MD and tMD OCC → ‘collapsed IF’ did not overlap well with each other (see figure S11bc), made us suspect that hysteresis is at play here. A comparison to the MEMENTO 2D-PMF is also suggestive of strong hysteresis, if we assume the MEMENTO results as ground truth. We cannot explicitly show the starting-state bias by contrasting PMFs from tMD in opposite directions, since we were unable to perform tMD in the IF → OCC direction at all. This in itself can be taken as strong indication for hysteresis, nonetheless.

We have demonstrated in conclusion that MEMENTO — together with an iterative approach for deriving CVs to match DOFs sampled in long unbiased MD — can provide valuable insight into complex and global protein conformational changes in LeuT. Targeted MD, in turn, fails again to provide paths of matching quality for the purposes of US, due to starting state bias.

## Discussion

We investigated in this work four protein systems that exhibit conformational changes for which structural data is available at both end states: deca-alanine, ADK, P38*α* and LeuT. Furthermore, for each of these systems, previous studies that calculated the PMFs for the conformational changes are available in the literature. The aim of this project was to determine the extent to which tMD suffers from hysteresis when used to seed umbrella sampling simulations for computing PMFs. We also wished to study whether these adverse effects can be mitigated by the use of paths generated with MEMENTO (implemented and freely available as PyMEMENTO) — a simple procedure we devised for crafting history-independent paths using a combination of coordinate morphing and template-based structure modelling.

On deca-alanine, we found no difference in the quality of tMD and MEMENTO paths: the simple nature of the system evidently does produce significant hysteresis. The example therefore serves as validation of the PyMEMENTO code, showing that it does not introduce artifacts in an easy test case. Moving on to ADK, tMD and MEMENTO paths can both yield converged PMFs along physically intuitive CVs. When attempting to investigate the role of an alternative closed conformation, however, hysteresis appeared in bi-directional tMD, thus conferring an advantage to MEMENTO for a thorough exploration of the system. For P38*α*, tMD suffered from large-scale hysteresis. MEMENTO, in turn, was able to generate converged and consistent PMFs, albeit only once we had derived custom CVs using an iterative scheme that utilises MEMENTO, extensive end-state sampling and PCA. Lastly, concerning the membrane transporter LeuT this approach for generating custom CVs was again successful with MEMENTO, while tMD suffered from hysteresis even when using the consistent CV space obtained from MEMENTO results. We note that for all examples, our results largely agree with previously published data in the literature.

In conclusion, tMD is prone to hysteresis when combined with umbrella sampling for all but the simplest conformational changes, at least without substantial prior knowledge regarding CV choice. We therefore suggest as best practise to all investigators who use tMD to generate paths for refinement and PMF computations to carefully use controls in both targeting directions, where possible. Hysteresis — if it is an issue within the relevant CV space — will become apparent as substantial differences between the obtained PMFs. It is unclear still whether this is also an issue when using tMD as a part of more elaborate biasing schemes, however our results warrant a degree of caution. It is left for future work to establish this for each of the many methods using tMD. We also show that MEMENTO, despite its conceptual simplicity and easy implementation, is a powerful and flexible tool for conformational sampling. This is especially the case due to the ease of its combination with long end-state sampling and dimensionality reduction for CV derivation, and because newly discovered conformational states can effortlessly be integrated into existing ensembles of paths. Nonetheless, the free energy of protein conformational changes remains a formidable challenge that requires substantial expertise and system-specific knowledge or deep *ad-hoc* investigation on the part of the researcher.

## Supporting information

Zipped archive of movies

## Acknowledgement

We thank Dr. Zhiyi Wu for training and helpful discussions during the early stages of the project, Dr. Irfan Alibay and Dr. Rocco Meli for their help with python software development, as well as all members of the Biggin group for their continued feedback. This project was funded by the Wellcome Trust (Grant ID: 218514/Z/19/Z) and compute resources at the EPSRC ARCHER2 and N8 CIR BEDE facilities, granted via the High-End Computing Consortium for Biomolecular Simulation (HECBioSim — https://www.hecbiosim.ac.uk/), supported by EPSRC (EP/L000253).

## Supporting Information Available

- Supplementary figures S1–S11. Supplied as one pdf file.
- A set of 15 supplementary videos illustrating the progresion of MEMENTO paths as discrete intermediates, as well as the collective variables we derived using principal component analysis as rocking animations. Supplied as separate mp4 and mpg files.
- Supplementary data with key coordinate files and simulation biasing and analysis output (in PLUMED format, and as PMFs produced by WHAM) is available at https://doi.org/10.5281/zenodo.7567883.

