## Supplementary Information for "Tackling hysteresis in conformational sampling — how to be forgetful with MEMENTO"

### Supplementary figures

This file contains 11 supplementary figures, as referenced in the main text.

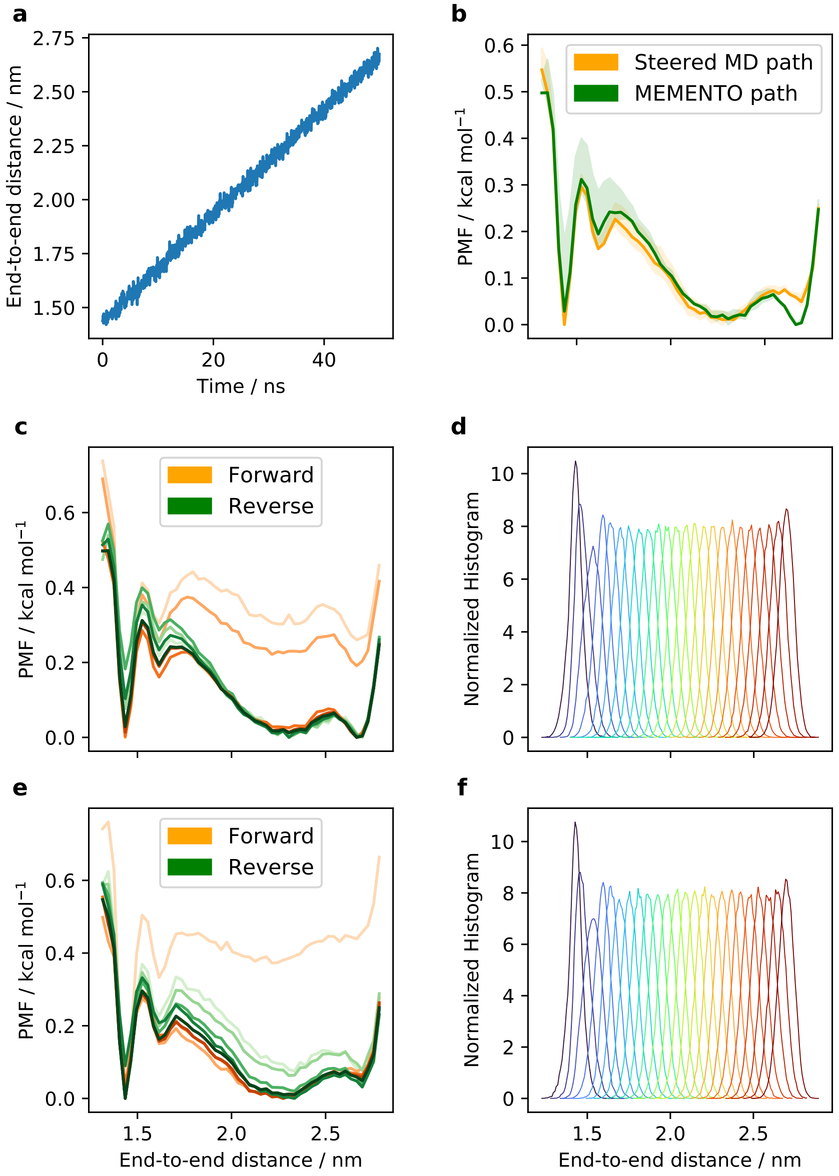


Figure S1: (a) Steered MD progression of deca-alanine along the end-to-end distance CV. (b) PMFs from 1D-REUS with steered MD and MEMENTO paths overlayed, displaying nearly identical shapes. Shaded area is the range of PMFs observed when taking only the first 60%, the last 60% and the full sampling. (c) Convergence of MEMENTO 1D-REUS, shown by the PMF taking (with increasing line saturation) 20, 40, 60, 80 and 100 % of sampling from the front (orange) or the back (green) of trajectories. (d) Normalised MEMENTO 1D-REUS histograms, showing excellent overlap. (e)–(f) are equivalent to (c)–(d), but show 1D-REUS derived from the steered MD path.

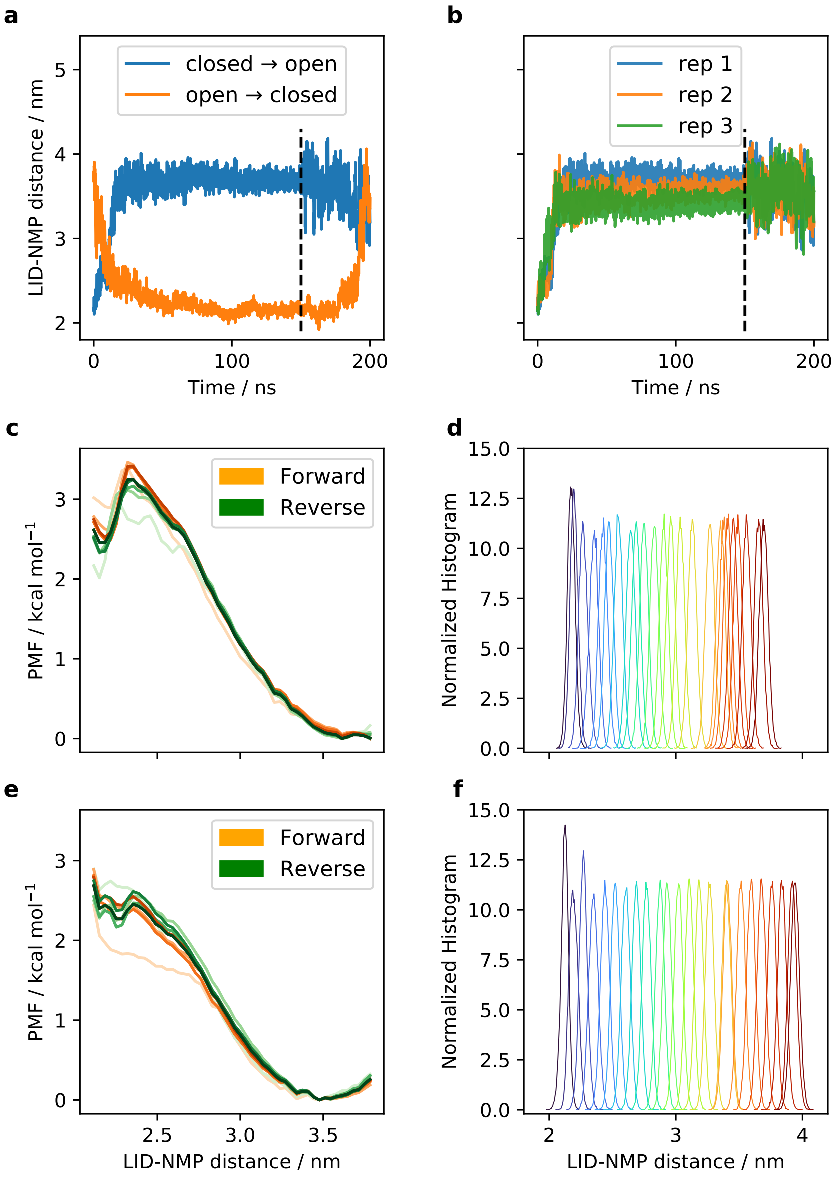


Figure S2: (a) ADK tMD based along the C$\alpha$-RMSD, projected onto the LID–NMP distance. The obtained closed conformation is not stable. (b) Three replicates of closed $\rightarrow$ open tMD all give stable open states. (c) Convergence of MEMENTO 1D-REUS rep 1, shown by the PMF taking (with increasing line saturation) 20, 40, 60, 80 and 100 % of sampling from the front (orange) or the back (green) of trajectories. (d) Normalised MEMENTO 1D-REUS rep 1 histograms, showing sufficient overlap. (e)–(f) are equivalent to (c)–(d), but show 1D-REUS rep 1 derived from a tMD path.

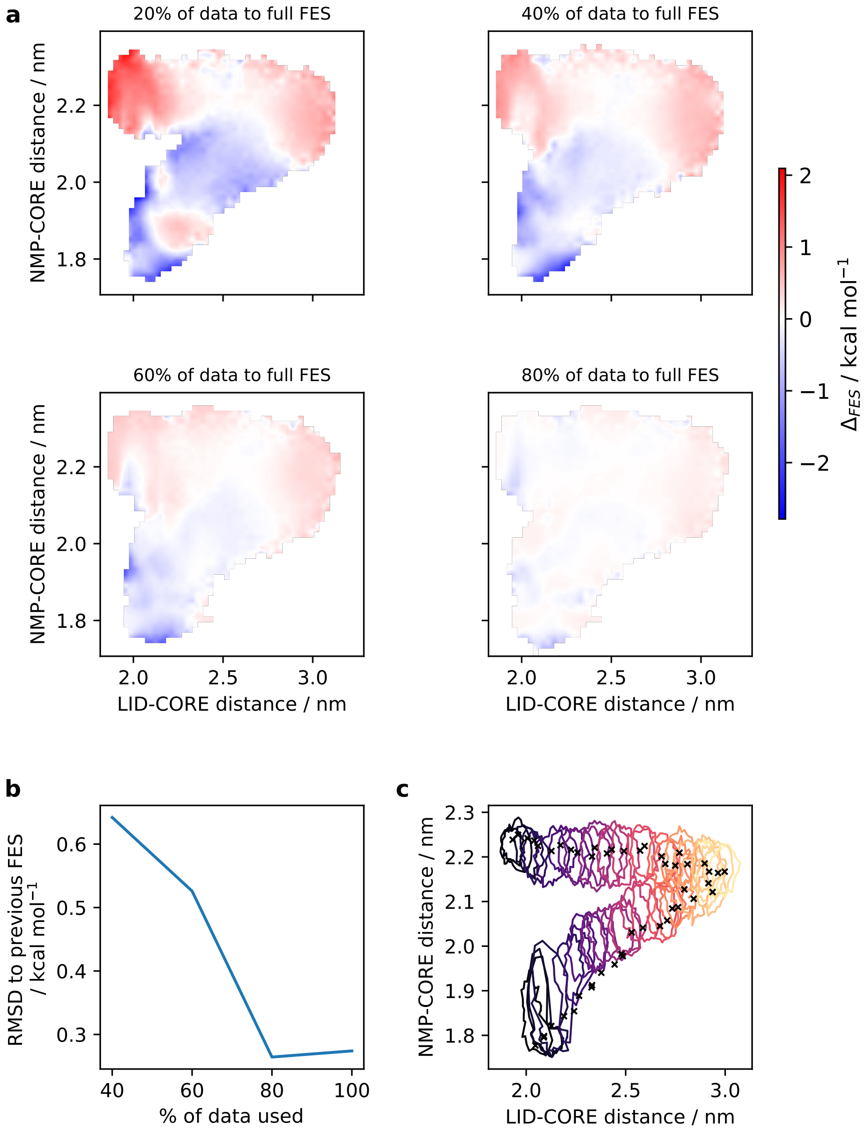


Figure S3: (a) Convergence of ADK 2D-REUS from MEMENTO paths, shown as the difference between the PMFs including 20, 40, 60 and 80% of the data and the full PMF, thus highlighting regions of faster and slower convergence. (b) Alternative representation of convergence as the RMSD from adding successive 20% chunks of data into the PMF computation. (c) 2D-histograms shown as paths encircling the normalised histograms at 30% of their peak height, displaying satisfactory overlap. Crosses indicate the REUS window starting frames.

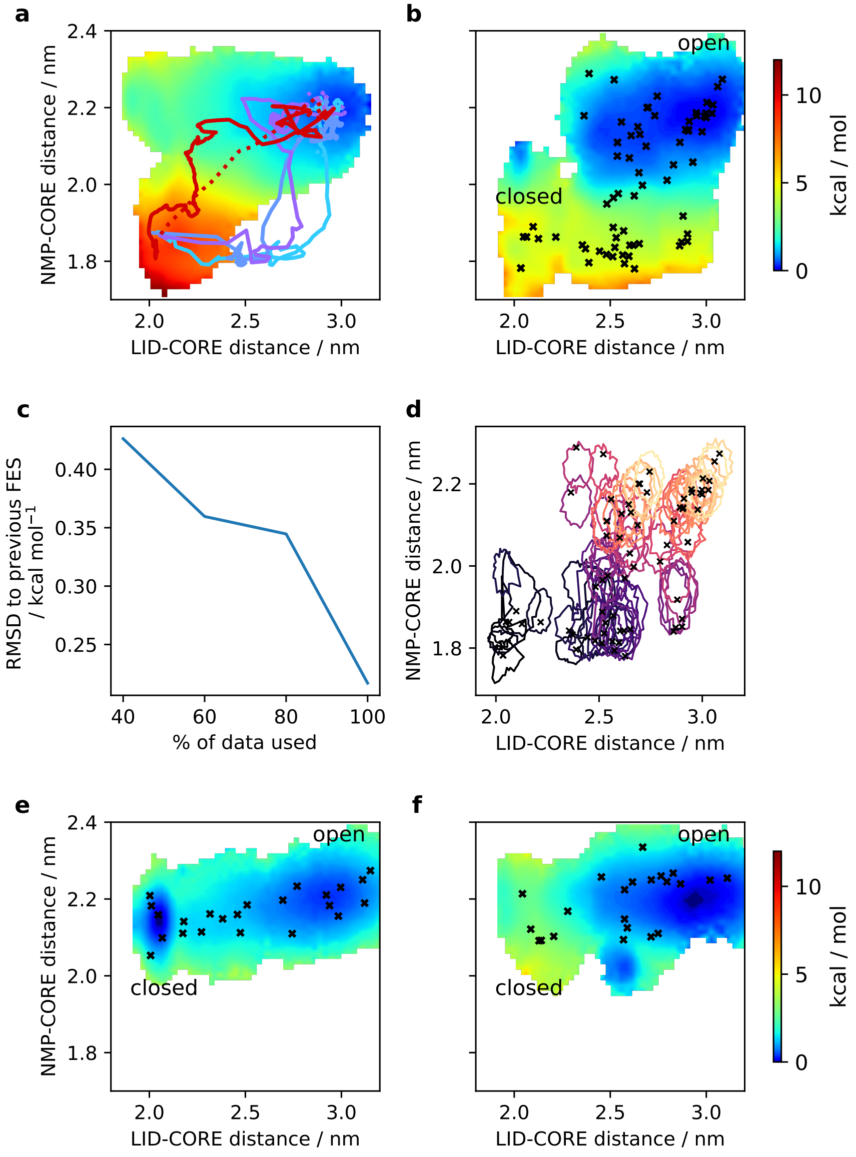


Figure S4: (a) Projection of the ADK tMD runs onto the LID–CORE and NMP–CORE distance 2D-CV space. Blue hues represent closed → open runs, the red line is the open → closed run. Solid lines are the biased portion, dashed lines the relaxing portion of the runs. The 2D-PMF from MEMENTO is shown as reference in the background. (b) PMF from 2D-REUS based on closed → open tMD runs. Crosses indicate the REUS window starting frames. (c) Convergence of the 2D-PMF as the RMSD from adding successive 20% chunks of data into the PMF computation. (d) 2D-histograms shown as paths encircling the normalised histograms at 30% of their peak height, overall satisfactory overlap with issues in some regions. (e)–(f) PMFs from 2D-REUS from the (e) alternative closed → open and (f) open → alternative closed tMD runs, showing significant bias towards the tMD starting state (hysteresis).

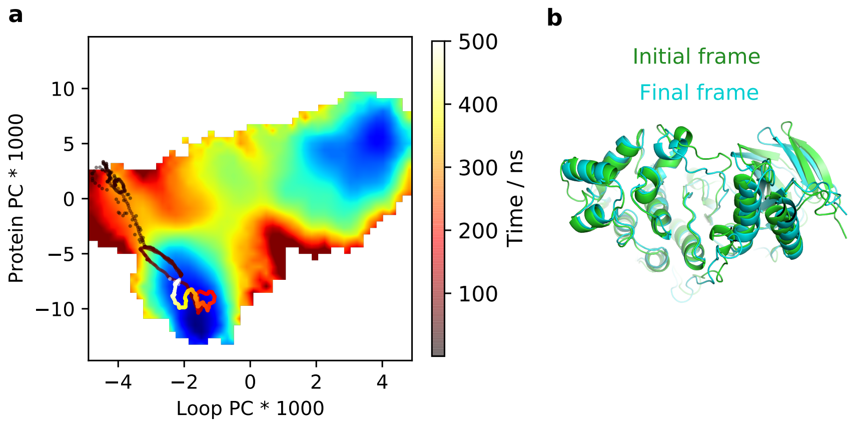


Figure S5: (a) Projection of the apo-P38$\alpha$ unbiased MD run starting from DFG-out onto the 2D-PMF with PCA-CVs, showing a relaxation towards an alternative DFG-out state. (b) Rendering of the alternative DFG-out state, showing a lesser extent of DFG-loop opening and a general twist in the protein.

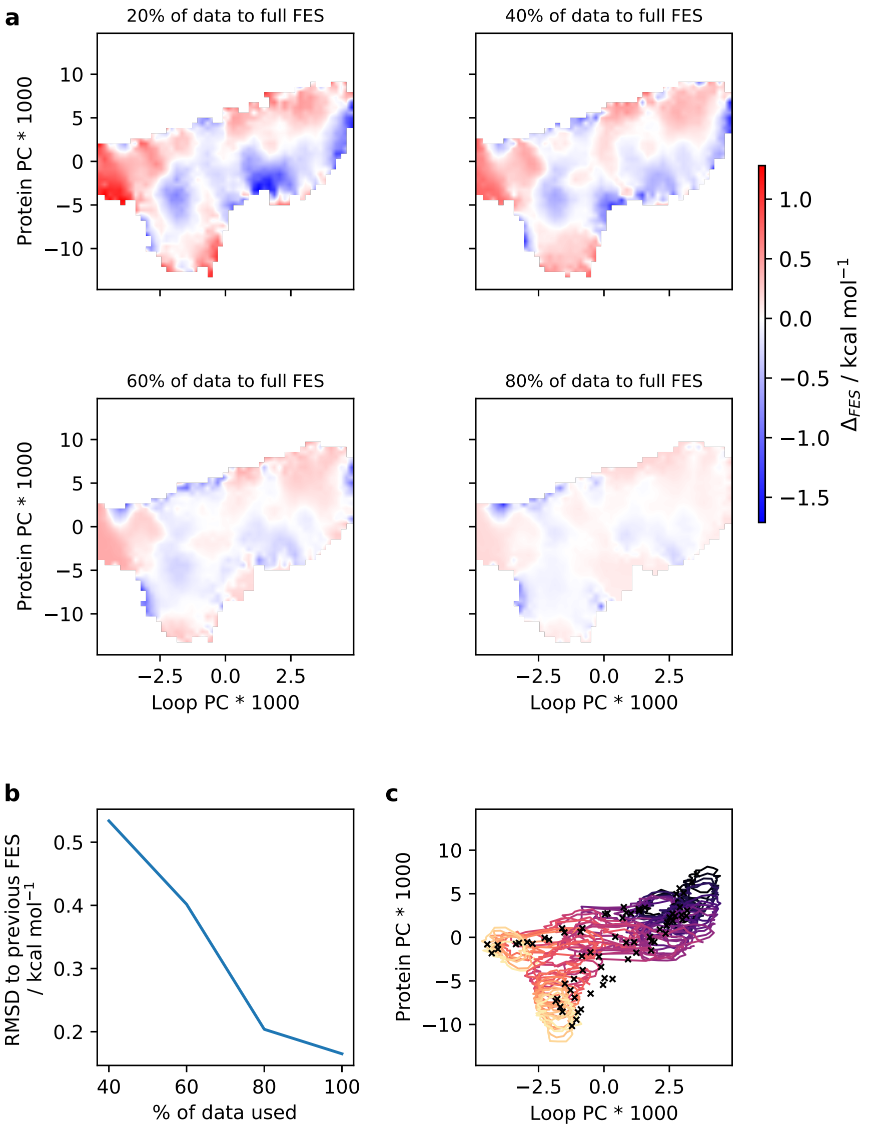


Figure S6: (a) Convergence of P38$\alpha$ 2D-REUS from MEMENTO paths, shown as the difference between the PMFs including 20, 40, 60 and 80% of the data and the full PMF, thus highlighting regions of faster and slower convergence. (b) Alternative representation of convergence as the RMSD from adding successive 20% chunks of data into the PMF computation. (c) 2D-histograms shown as paths encircling the normalised histograms at 30% of their peak height, displaying satisfactory overlap. Crosses indicate the REUS window starting frames.

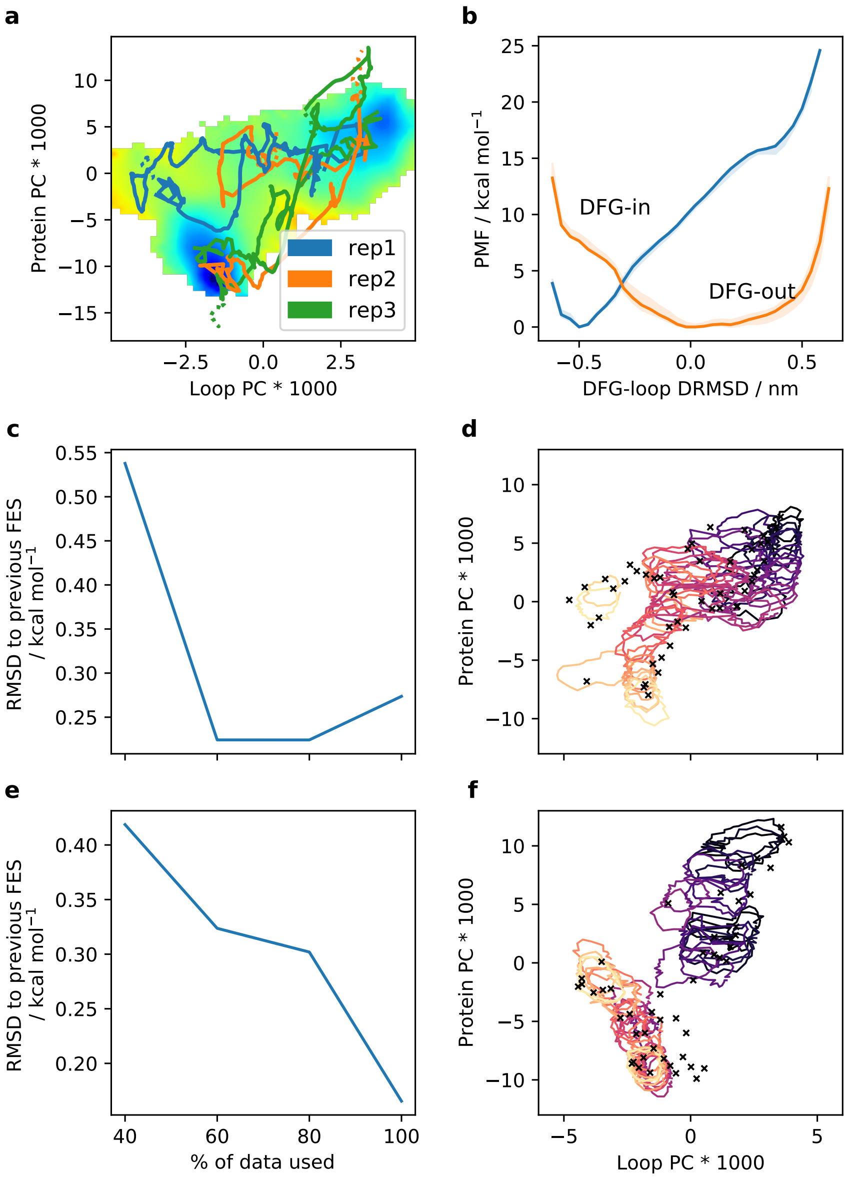


Figure S7: (a) Projection of the P38$\alpha$ tMD runs onto the PCA CV-space. Solid lines are the biased portion, dashed lines the relaxing portion of the runs. The 2D-PMF from MEMENTO is shown as reference in the background. Rep 1 and 3 approach their respective target states in both directions, while the rep 2 tMD fails in the DFG-in $\rightarrow$ DFG-out direction. (b) 1D-REUS from tMD rep 1 paths after re-solvation of all intermediates. (c) Convergence of the 2D-PMF for DFG-in $\rightarrow$ DFG-out tMD, as the RMSD from adding successive 20% chunks of data into the PMF computation. (d) 2D-histograms for the same PMF, shown as paths encircling the normalised histograms at 30% of their peak height, with some overlap issues. (e)–(f) Same as (c)–(d), but for the DFG-out $\rightarrow$ DFG-in tMD direction.

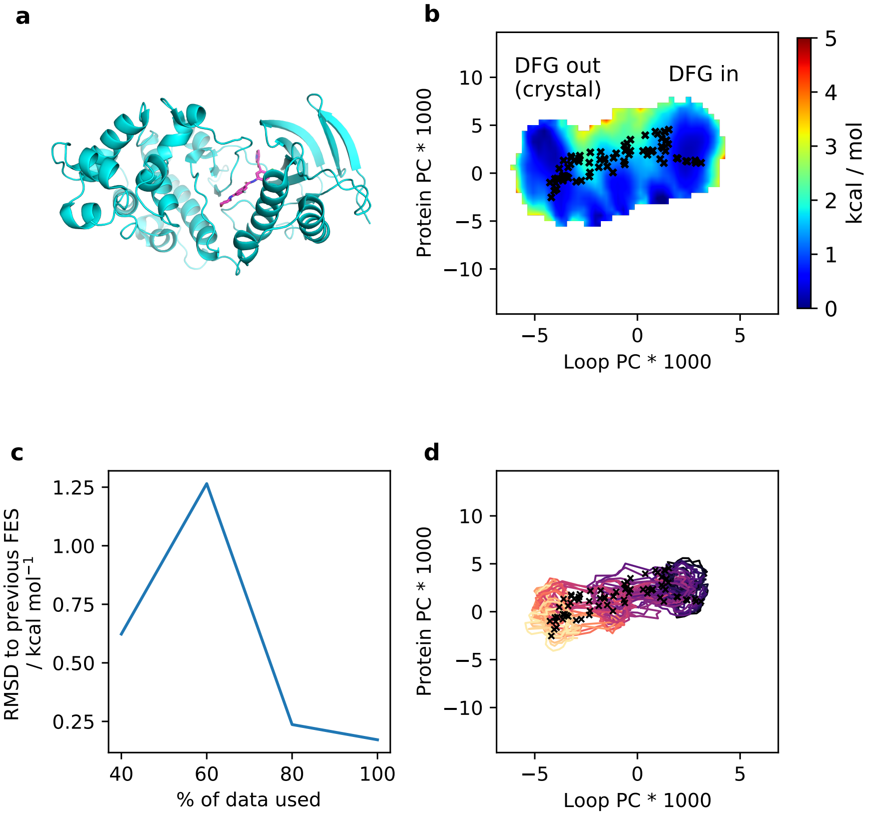


Figure S8: (a) P38$\alpha$ inhibitor binding pose in the DFG-out conformation. (b) 2D-REUS along PCA-CVs with MEMENTO paths for holo P38$\alpha$. Crosses indicate the REUS window starting frames. (c) Convergence of the 2D-PMF as the RMSD from adding successive 20% chunks of data into the PMF computation. (d) 2D-histograms, shown as paths encircling the normalised histograms at 30% of their peak height, with satisfactory overlap

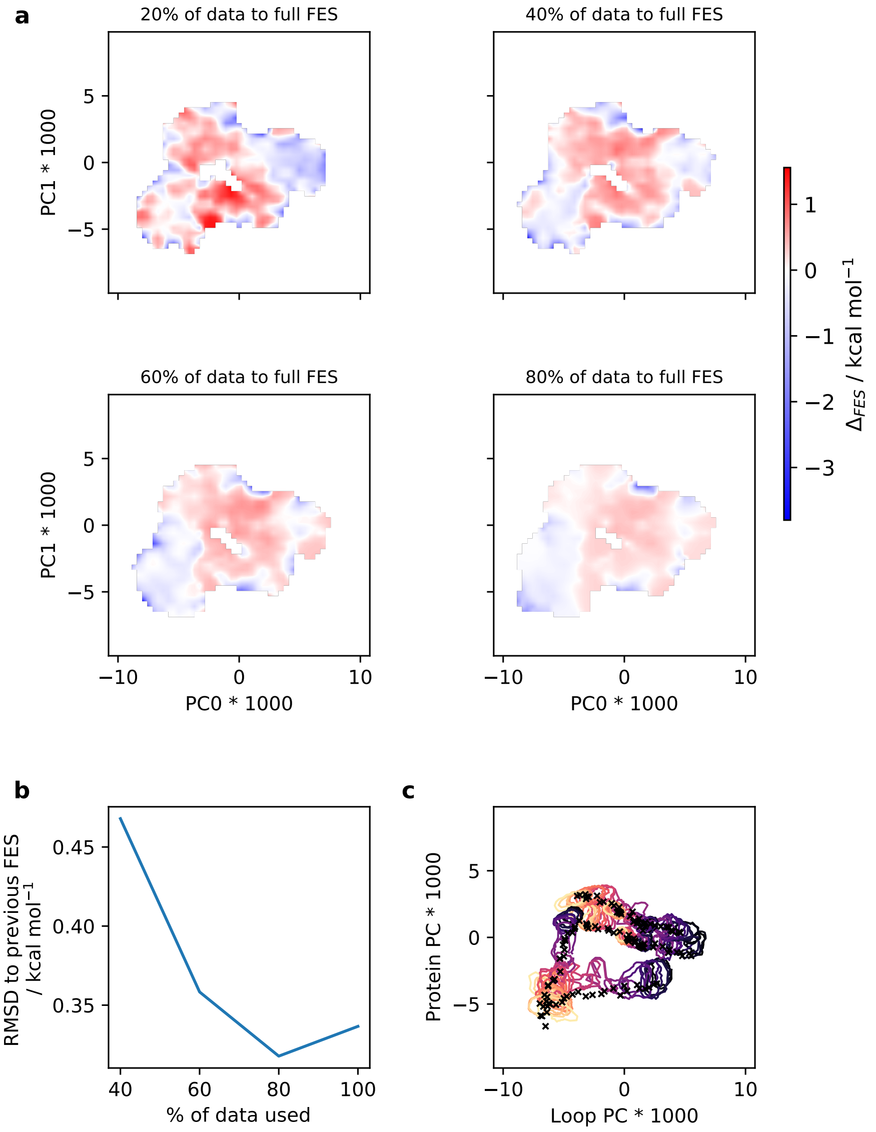


Figure S9: (a) Convergence of LeuT 2D-REUS from MEMENTO paths, shown as the difference between the PMFs including 20, 40, 60 and 80% of the data and the full PMF, thus highlighting regions of faster and slower convergence. (b) Alternative representation of convergence as the RMSD from adding successive 20% chunks of data into the PMF computation. (c) 2D-histograms shown as paths encircling the normalised histograms at 30% of their peak height, displaying satisfactory overlap. Crosses indicate the REUS window starting frames.

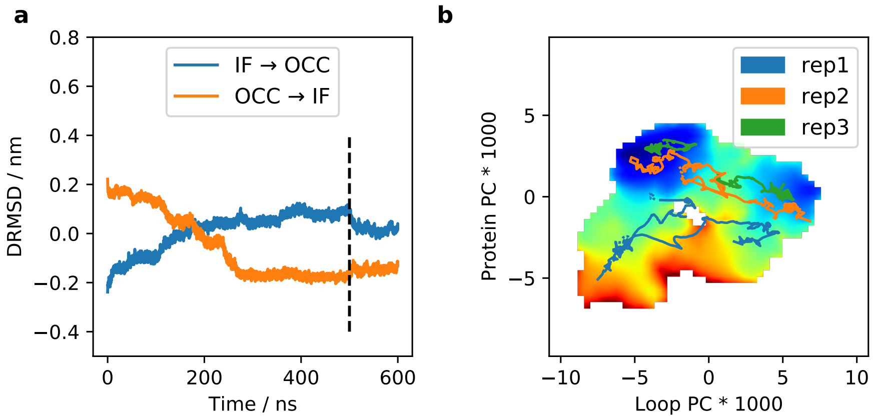


Figure S10: (a) LeuT tMD between IF and OCC states, replicate 1, projected onto the DMRSD CV. The dashed line represents the end of the restraint, followed by unbiased relaxation. (b) Three replicates of LeuT tMD, projected onto the MEMENTO 2D-PMF, showing how only replicates 1–2 in the OCC → IF direction reached their respective target states. Solid lines are the biased portion, dashed lines the relaxing portion of the runs.

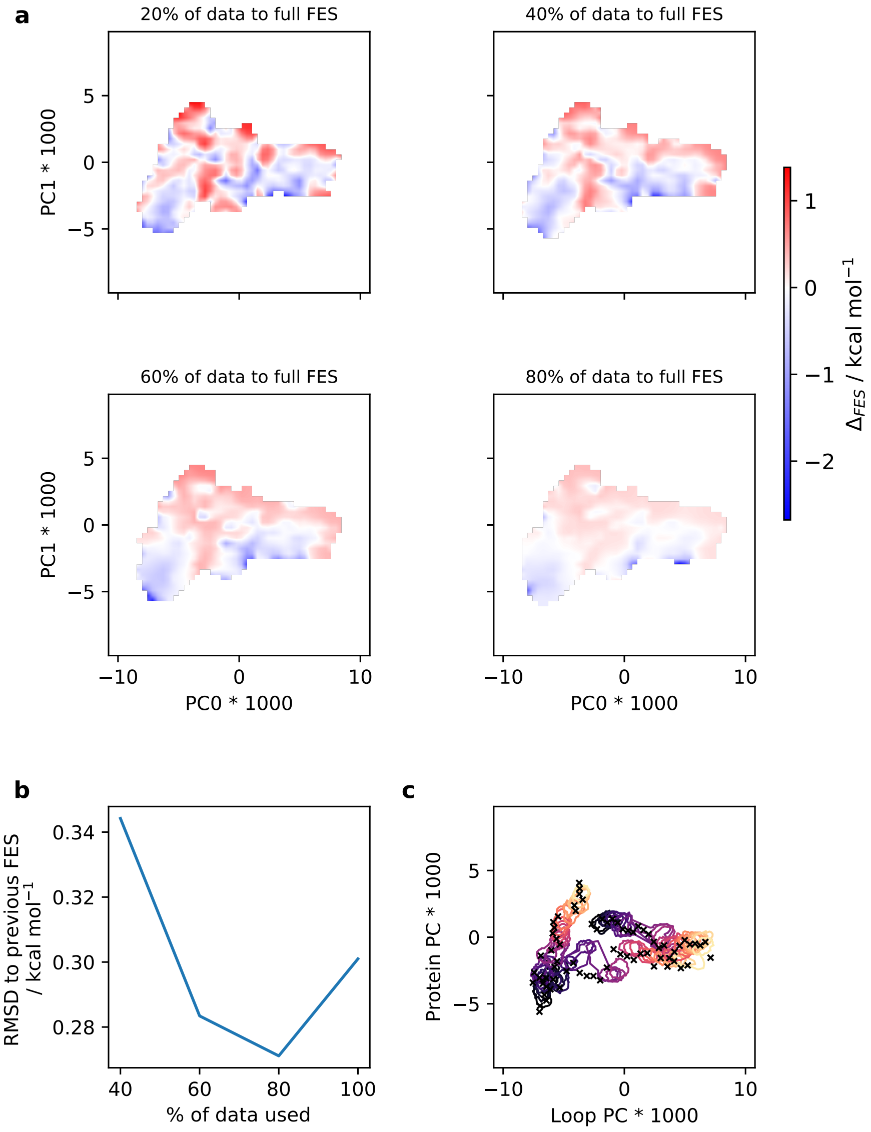


Figure S11: (a) Convergence of LeuT 2D-REUS from tMD paths, shown as the difference between the PMFs including 20, 40, 60 and 80% of the data and the full PMF, thus highlighting regions of faster and slower convergence. (b) Alternative representation of convergence as the RMSD from adding successive 20% chunks of data into the PMF computation. (c) 2D-histograms shown as paths encircling the normalised histograms at 30% of their peak height, displaying insufficient overlap. Crosses indicate the REUS window starting frames.
